# Rasputin/G3BP mediates o’nyong-nyong virus subversion of antiviral immunity in Anopheles coluzzii

**DOI:** 10.1101/2022.11.23.517724

**Authors:** Solène Cottis, Adrien Blisnick, Christian Mitri, Emma Brito-Fravallo, Mariette Matondo, Anna-Bella Failloux, Kenneth D. Vernick

**Affiliations:** Institut Pasteur, Université de Paris, CNRS UMR2000, Genetics and Genomics of Insect Vectors Unit, Department of Parasites and Insect Vectors, F-75015 Paris, France; Graduate School of Life Sciences ED515, Sorbonne Universités UPMC Paris VI, 75252 Paris, France; Institut Pasteur, Université de Paris, Arboviruses and Insect Vectors Unit, Department of Virology, F-75015 Paris, France; Institut Pasteur, Université de Paris, Mass Spectrometry for Biology Platform UTECHS, CNRS UAR2024, Department of Structural Biology and Chemistry, F-75015 Paris, France

**Author notes:** Correspondence to KDV. Email addresses SC.

**Keywords:** Arbovirus, G3BP, host-pathogen interaction, innate immunity

## Abstract

The G3BP proteins in vertebrates and Aedes mosquito ortholog, Rasputin, are essential for alphavirus infection, but the underlying mechanism of Rasputin/G3BP proviral activity is poorly understood. It has been suggested that G3BP could influence host immune signaling, but this has not been functionally demonstrated. Here, we find that depletion of Rasputin activity in Anopheles mosquitoes, the primary vectors of the alphavirus o’nyong-nyong (ONNV), provokes dysregulation of the antiviral Imd, JAK/STAT and RNAi pathways, indicating that Rasputin is required for expression of normal basal immunity in uninfected mosquitoes. Depletion of Rasputin during ONNV bloodmeal infection causes increased transcript abundance of genes in the Imd pathway including positive regulator Rel2, and decreases ONNV infection in mosquitoes. Loss of Rasputin is complemented by co-depletion of Imd pathway positive regulator, Rel2, which restores normal ONNV infection levels. Thus, the presence of Rasputin is required for ONNV inhibition of Imd activity, and viral inhibition of Imd explains much of the Rasputin proviral activity. The viral non-structural protein 3 (nsP3) binds to Rasputin and alters the profile of cellular proteins binding to Rasputin. In the presence of nsP3, 48 Rasputin-binding proteins are unchanged but seven binding proteins are excluded and eight new proteins bind Rasputin. The Rasputin binding partners altered by nsP3 are candidate factors for ONNV immune manipulation and subversion through Rasputin. Overall, these results are consistent with and strongly suggest a mechanism in which ONNV, probably nsP3, co-opts the normal Rasputin function assuring basal cellular immune activity in order to inhibit antiviral immunity and promote infection. These observations may be generalizable for Rasputin function during alphavirus infection of other mosquitoes, as well as for G3BP function in the mammalian host, and could offer a target for vector-based control of arbovirus transmission.

## INTRODUCTION

Arthropod-borne viruses (arboviruses) are maintained through an alternating cycle of transmission between mammalian hosts and arthropod vectors. Arboviruses represent a spreading global burden for human and animal health, with the clinically most important pathogens represented by RNA viruses in the families Flaviviridae, Togaviridae, Bunyavirales, and Reoviridae (Weaver and Reisen, 2010). The alphavirus o’nyong-nyong (ONNV, genus alphavirus, family Togaviridae) is phylogenetically closely related to chikungunya virus (CHIKV, genus alphavirus, family Togaviridae). Both are in the same antigenic group, the Semliki forest virus complex, thus are difficult to distinguish by immunodiagnostic assay, and symptomatically cause essentially the same human disease (Powers et al., 2001; Rezza et al., 2017). The most apparent difference between the arboviruses is their use of mosquito vectors. Anopheles mosquitoes are the major vector of human malaria but are the primary vector of only one known arbovirus, ONNV, while Aedes mosquitoes transmit the majority of mosquito-borne arboviruses, including CHIKV.

The molecular mechanism underlying the mosquito vector specificity of ONNV and CHIKV is not understood. Evidence points to the involvement of non-structural protein 3 (nsP3), which is diverged between the two viruses. Replacing CHIKV nsp3 with the ONNV nsP3 gene in the CHIKV genomic backbone allowed partial infection of Anopheles mosquitoes, while the CHIKV backbone with its own nsP3 gene is noninfective to Anopheles (Saxton-Shaw et al., 2013; Vanlandingham et al., 2005; Vanlandingham et al., 2006). Interestingly, unlike CHIKV, which does not infect Anopheles, ONNV can infect some tested strains of Ae. aegypti (Vanlandingham *et al*., 2005). These results suggest that Anopheles is the more stringent or less permissive host among the two for virus infection, and also imply that observed viral nsP3 sequence divergence may be based on specific viral adaptation to infect either Anopheles or Aedes hosts, because the human host is shared by both viruses.

The probable role of viral nsP3 in vector host specificity points in turn to the a host protein known to bind nsP3 of many alphaviruses, the Ras-GTPase-activating protein (SH3 domain)-binding protein (G3BP) and the mosquito ortholog Rasputin (Rin) (Panas et al., 2014). Rin/G3BPs are enigmatic molecules with diverse biological functions (Alam and Kennedy, 2019; Kang et al., 2021). Most studies of Rin/G3BP function have been carried out in mammalian cells, where there are three isoforms (G3BP1, G3BP2a and 2b) expressed from two genes (Irvine et al., 2004; Kang *et al*., 2021). Mosquitoes carry a single Rin gene, with the small number of published studies restricted to Aedes mosquitoes (Fros et al., 2015; Goertz et al., 2018; Nowee et al., 2021), and to our knowledge there are no previous published reports about Rin in Anopheles mosquitoes.

Mammalian G3BPs are involved in numerous biological processes including RNA-binding, RNA metabolism, stress granule formation, and regulation of ubiquitin-mediated degradation signaling (Alam and Kennedy, 2019; Irvine *et al*., 2004; Kang *et al*., 2021; Kedersha et al., 2016; Laver et al., 2020; Soncini et al., 2001). Aedes Rin can interact and colocalize with nsP3 of many tested alphaviruses in Aedes cells (Fros *et al*., 2015; Goertz *et al*., 2018; Nowee *et al*., 2021). Rin/G3BPs also serve as proviral host factors promoting the infection cycle of multiple RNA viruses in mammals, while in Aedes proviral activity has been demonstrated only for alphaviruses (Fros *et al*., 2015). The proviral function of Rin/G3BPs is associated in general with efficiency of viral genome replication, although “the exact mechanism by which they act remains to be explored” (Scholte et al., 2015). Interestingly, G3BPs have also been reported to potentially interact with host immunity by influencing cytoplasmic distribution of the regulators of the NF-kappa B pathway (Prigent et al., 2000), by enhancing NF-kappa B phosphorylation and protein quantity (Scholte *et al*., 2015; Zhang et al., 2012), and by modulating NF-kappa B signaling through protein kinase R or by other means (Deater et al., 2022; Reineke and Lloyd, 2015). The implications of potential Rin/G3BP interaction with immune signaling factors upon actual immune regulation, or on the efficiency if virus infection have barely been examined, and there are no studies in mosquitoes.

Because the above evidence points to differential interactions of viral nsp3 with host Rin to explain at least in part the observed vector specificity of ONNV and CHIKV for infection of Anopheles and Aedes, here we undertook a study of the function of Anopheles Rin in ONNV infection, which is a knowledge gap. More broadly, the reason for the apparent paucity of arbovirus transmission by Anopheles mosquitoes in nature is also not understood, because the study of virus-host interactions in Anopheles has been relatively neglected in favor of the rich extensive research on Anopheles-Plasmodium interactions. However, Anopheles harbor a complex natural virome of RNA viruses (Belda et al., 2019) and a number of pathogenic arboviruses have been isolated from Anopheles mosquitoes (Nanfack Minkeu and Vernick, 2018). For example, there is evidence of Anopheles contribution to maintenance of Rift Valley Fever virus during epidemics (RVFV, genus phlebovirus, family Bunyavirales) (Ratovonjato et al., 2011; Seufi and Galal, 2010). These observations present a biological puzzle that highlights the importance of studying Anopheles-virus interactions. With changing climatic and ecological conditions and exposure of vector populations to new potential pathogens in new environments, it is relevant to ask what it would require for arboviruses other than ONNV to adapt to Anopheles mosquitoes as transmission vectors.

Here we investigated the proviral activity of Anopheles Rin, and discovered a clear and direct link between Rin and antiviral immunity. We found that Rin influences antiviral immunity in ONNV infection of Anopheles. and probably underlies viral co-optation by nsP3 of host Rin in order to subvert antiviral immunity. The findings could be relevant to studies of G3BPs in mammals and for other alphaviruses.

## RESULTS

### Rin activity is required for normal basal expression of multiple immune pathways in uninfected Anopheles

Rin/G3BP is proviral for infection of many alphaviruses in mammalian cells and for infection of CHIKV in Aedes albopictus (Fros *et al*., 2015; Gotte et al., 2020; Scholte *et al*., 2015) but it has not been examined in Anopheles. Therefore, we first, confirmed that Rin is proviral for ONNV in Anopheles coluzzii mosquitoes by depleting Rin transcript using RNAi-mediated gene silencing by injection of double-stranded RNA (dsRNA) specific for Rin (dsRin), or irrelevant GFP control (dsGFP), followed by bloodmeal infection with ONNV. The depletion of Rin in An. coluzzii significantly decreased ONNV infection prevalence (defined as the proportion of fully fed mosquitoes positive for ONNV) and infection intensity (defined as the viral titer measured in the ONNV-positive mosquitoes) (Fig. S1A and S1B). Therefore, Rin activity is proviral and agonistic for ONNV infection in An. coluzzii by infectious bloodmeal.

Because of the suggestive evidence from mammalian G3BPs detailed above, we hypothesized that Rin proviral activity could be explained, at least in part, if Rin is required for host immune signaling, and if a viral virulence factor can manipulate Rin to undermine cellular immune signaling in relevant antiviral pathways. Therefore, we first determined whether Rin influences basal immunity in uninfected mosquitoes by measuring the influence of Rin depletion by dsRin treatment as compared to dsGFP controls upon transcript levels of a panel of immune genes selected on two main lines of evidence: previous functional demonstration of antiviral activity against ONNV in An. coluzzii (Carissimo et al., 2015), and differential RNAseq transcript abundance in An. coluzzii after ONNV bloodmeal infection (Carissimo et al., 2018). The candidate panel is comprised of positive regulators of the four main mosquito immune pathways (Toll, Imd, JAK/STAT and small RNA), and immune factors or effectors, including the leucine-rich repeat protein (LRR) genes APL1C, APL1A, LRIM1, LRIM4, LRIM10; the complement-like thioester proteins TEP3, TEP4, TEP12; and inducer of immune granulocyte differentiation, Evokin).

First, we queried Rin influence on expression of Toll pathway regulatory genes (positive regulator, Rel1 and negative regulator, Cactus) and effector genes (APL1C and TEP3) (Fig. 1A). APL1C and TEP3 are extracellular immune factors regulated by the Toll pathway (Mitri et al., 2015), and APL1C displays antiviral activity against ONNV in An. coluzzii, while TEP3 does not (Carissimo *et al*., 2015). Depletion of Rin has little influence on Toll-pathway related immune transcripts except for causing a late increase of APL1C transcript abundance, indicating that normal Rin activity slightly limits APL1C transcript abundance in uninfected mosquitoes.

**Figure 1.**
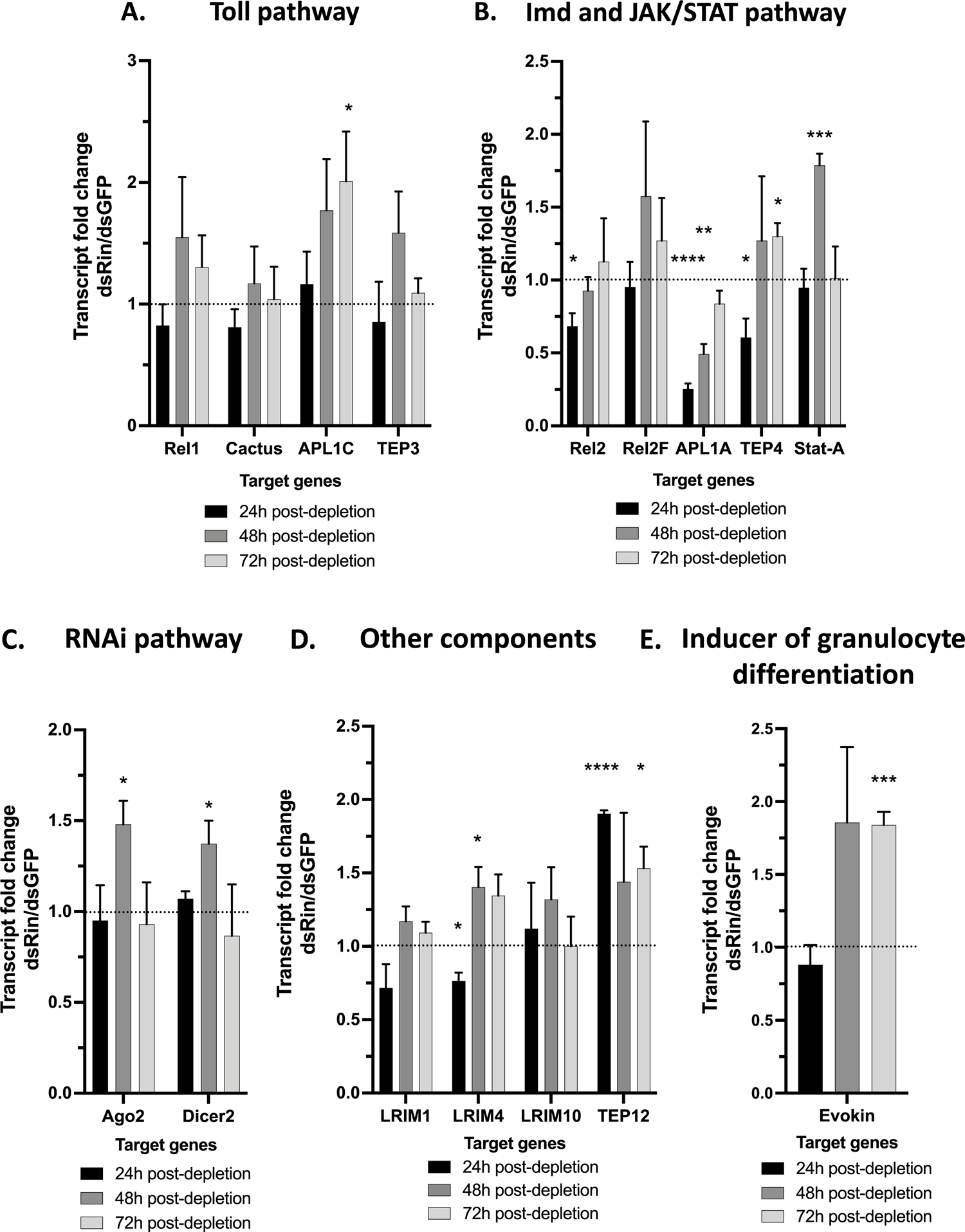
Rin function is required for correct transcriptional basal immunity in uninfected Anopheles mosquitoes. (A-F) Female mosquitoes were injected with 500 ng/µL of dsRin or dsGFP. The influence of Rin depletion on genes involved in the Toll pathway (A), the Imd pathway (B), the JAK/STAT pathway (B), the siRNA pathway (C), other immune transcripts involved in complement-like immunity including LRR and TEP transcript (D) and the lipid carrier Evokin (E) are showed. Each bar represents the level of transcript abundance of each immune genes relative to dsGFP-treated control (defined as 1.0), and error bars indicate the SEM. Histogram showed are the results of three independent experiments with 18 female mosquitoes analyzed in total per conditions. *P < 0.05, ** P < 0.01, *** P < 0.005, **** P < 0.0001.

In contrast, depletion of Rin in uninfected mosquitoes triggered a more striking effect upon the Imd and JAK/STAT pathways (Fig. 1B), which are both strongly antiviral for ONNV in Anopheles (Carissimo *et al*., 2015). In the Imd pathway, Rin depletion caused decreased transcript abundance of the Imd positive regulator Rel2 as well as the Imd effector genes, APL1A and TEP4, which are transcriptionally regulated by Rel2 (Mitri *et al*., 2015). In the JAK/STAT pathway, Rin depletion caused increased transcript abundance of the JAK/STAT positive regulator, Stat-A, and therefore Rin is required to limit Stat-A transcript to normal levels. Thus, depletion of Rin causes widespread dysregulation of the Imd and JAK/STAT pathways, and normal Rin activity is required for correctly regulated basal activity of these immune pathways.

Among the small RNA immune pathways, the siRNA pathway has been studied for its role in counteracting RNA virus infection, as shown for flaviviruses such as ZIKV (Magalhaes et al., 2019) and DENV (Olmo et al., 2018), and alphaviruses CHIKV (Dong et al., 2022) and ONNV (Carissimo *et al*., 2015). The endonucleases Argonaute-2 (Ago2) and Dicer-2 (Dcr2) are essential catalytic components of the RNA-induced silencing complex (Kumar et al., 2018), and Ago2 was previously shown to be antiviral for ONNV infection (Carissimo *et al*., 2015). The transcript abundance of Ago2 and Dicer2 are both increased after Rin depletion, and therefore Rin acts to limit the expression of the siRNA pathway in uninfected mosquitoes (Fig. 1C).

We analyzed additional important extracellular immune factors for their transcriptional response to Rin. The LRR genes LRIM4 and LRIM10 are upregulated and downregulated, respectively, during ONNV infection of An. coluzzii (Carissimo *et al*., 2018). Here, we found that LRIM4 but not LRIM10 transcript abundance is altered by Rin depletion (Fig. 1D). The complement-like effector TEP12 is strongly increased by Rin depletion (Fig. 1D). Finally, Evokin is an Anopheles positive regulator for the proliferation of immune-competent hemocytes (Ramirez et al., 2015), and here we found that Evokin transcript is induced by Rin depletion (Fig. 1E), meaning that normal Rin function probably modulates the level of immune-primed hemocytes.

Taken together, these results in uninfected mosquitoes indicate that Rin is deeply involved in the correct basal regulation of key components of all four of the major arms of mosquito innate immunity, but particularly Imd and JAK/STAT. Absence of Rin leads to dysregulated expression of factors in all pathways. Rin is thus required to maintain the overall integrity and homeostasis of the systems necessary for immune surveillance and readiness of mosquitoes in the unstimulated resting state. Multiple of the Rin-regulated factors are known to be antiviral for ONNV in Anopheles, thus strengthening the hypothesis that Rin proviral function could be linked to viral manipulation of host immunity.

### Presence of Rin is required for ONNV-dependent inhibition of the Anopheles Imd pathway

After confirming above that Rin is proviral for ONNV infection of Anopheles, and that Rin activity is essential for correct expression of basal immunity in uninfected mosquitoes, we next examine the hypothesis that the proviral activity of Rin and the G3BP family could at least in part be explained by viral immune manipulation targeting Rin. The influence of Rin on Anopheles immunity was queried after challenge with an ONNV infective bloodmeal. Unfed mosquitoes were removed after the bloodmeal, and only fully engorged females were included in the analyses. Transcript abundance of the immune gene panel was measured at 48 h and 72 h after the infective bloodmeal. At 3 d post-bloodmeal, the ONNV infection is still restricted to the midgut epithelium, the primary midgut infection stage (PMI), prior to midgut escape and establishment of the systemic disseminated infection (Carissimo *et al*., 2015).

Rin depletion during the PMI of ONNV increases the transcript levels of genes in three of the four major immune signaling pathways, as compared to transcript levels in mosquitoes treated with dsGFP and then infected (Fig. 2 A-D). However, the major impact was observed in the Imd pathway, where the Imd positive regulator Rel2, ONNV antiviral effector APL1A (Carissimo *et al*., 2015), and ONNV induced LRR factor LRIM4 (Carissimo *et al*., 2018) are all significantly inhibited by Rin depletion, meaning that Rin activity is required for the decrease of the genes during the ONNV primary infection (Fig. 2B). Interestingly, although Imd and JAK/STAT were the two immune pathways that displayed the strongest antiviral effect against ONNV in the Anopheles PMI (Carissimo *et al*., 2015), here we find that the highly antiviral positive regulator of JAK/STAT, Stat-A, remains unaffected by the loss of Rin during ONNV infection (Fig. 2B).

**Figure 2.**
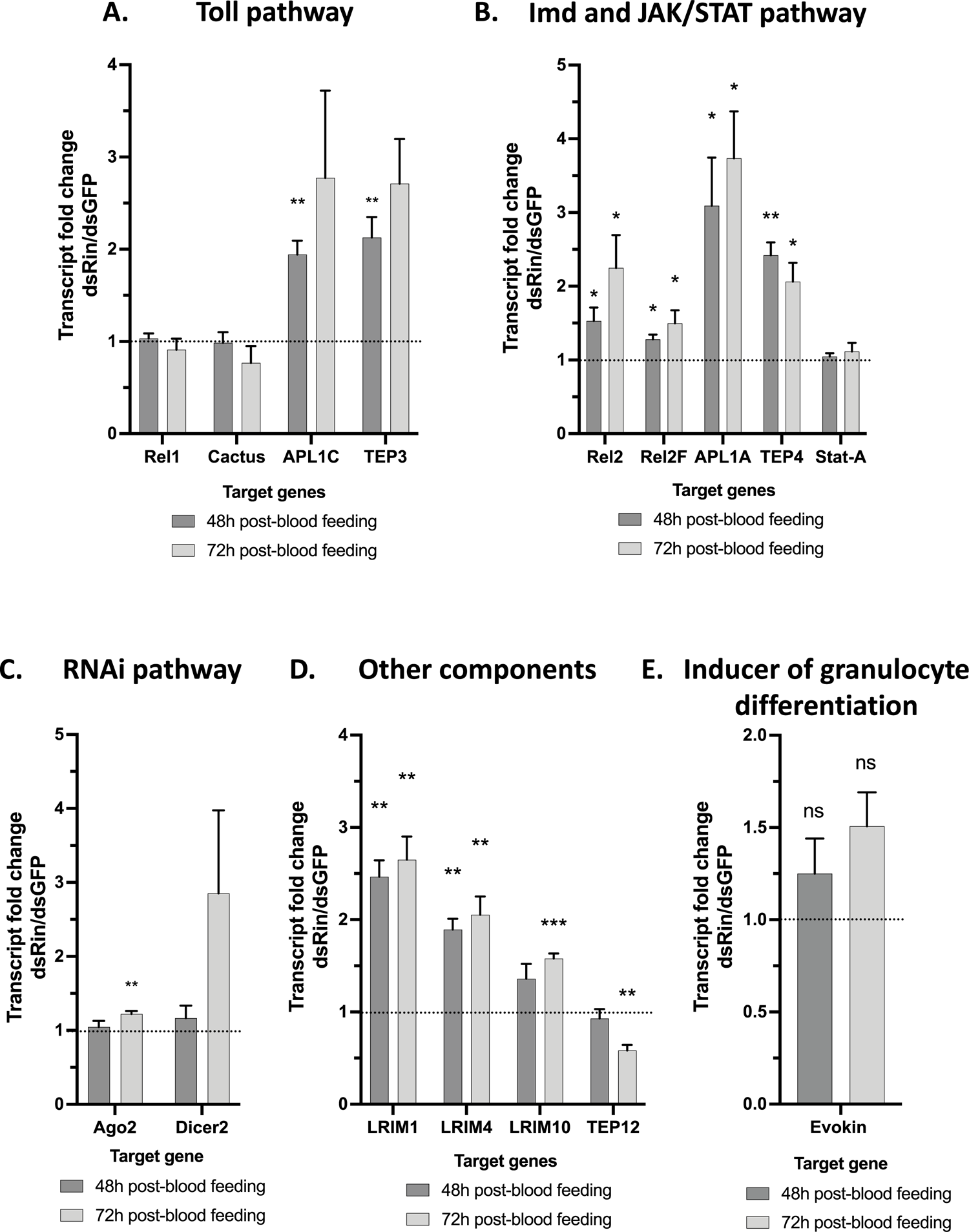
Rin is required for ONNV-dependent inhibition of the Anopheles Imd pathway. (A-E) Female mosquitoes were injected with 500 ng/µL of dsRin or dsGFP and two days later those females were fed an infectious blood meal containing ONNV at 10^7^ ffu/mL. Pull of 8 blood-fed mosquitoes were sacrificed at 2 days and 3 days post-blood feeding and total RNA was extracted. The influence of Rin depletion on genes involved in the Toll pathway (A), the Imd pathway (B), the JAK/STAT pathway (B), the siRNA pathway (C), LRR and TEP transcripts (D) and the lipid carrier Evokin (D) during the PMI are showed. Each bar represents the level of transcript abundance of each immune genes relative to dsGFP-treated control (defined as 1.0), and error bars indicate the SEM. Histogram showed are the results of three independent experiments with 48 female mosquitoes analyzed in total per conditions. *P < 0.05, ** P < 0.01, *** P < 0.005, ns non-significant.

There was only a small Rin influence on the Toll pathway during ONNV infection, where the positive and negative pathway regulators, Rel1 and Cactus respectively, are unchanged during the PMI by loss of Rin, although the Toll-regulated effector APL1C (Mitri et al., 2009), which is antiviral for ONNV (Carissimo *et al*., 2015), requires Rin for ONNV-dependent inhibition (Fig. 2A). In the siRNA pathway, Ago2, the critical “slicer” enzyme of the RNA-induced silencing complex (RISC), was slightly elevated 3 d after Rin depletion (Fig. 2C), indicating that there could be Rin-dependent inhibition of RNAi activity late in the PMI of ONNV. The RNAi pathway activity is not functionally antiviral for ONNV in the Anopheles PMI but only in the subsequent disseminated infection beginning after 3 d post-infection (Carissimo *et al*., 2015), and thus the late PMI effect of Rin on RNAi could indicate the beginning of viral midgut escape and the transition to the host antiviral response of the disseminated infection.

Above we showed that Rin is required to increase the Rel2 transcription factor to normal basal levels in uninfected mosquitoes (Fig. 1B), but during the PMI the presence of Rin (dsGFP treatment + ONNV infection) causes an ONNV-dependent decrease of Rel2 relative to the condition with depleted Rin and infection (Fig. 2B). Therefore, Rin blocks the induction of Rel2 that would be seen during ONNV infection in the absence of Rin (dsRin treatment + ONNV infection). The ability of Rin to modulate Rel2 expression could potentially be exploited by ONNV during the PMI to reduce Imd pathway activation, and therefore leads to the hypothesis that Rin could be a target for viral manipulation in order to inhibit the potent antiviral Imd pathway.

### Rin proviral activity is based on ONNV-dependent inhibition of the Imd pathway

Based on the above observation that the effect of Rin during ONNV infection is most strongly directed towards Imd as compared to the other immune pathways, we inferred that the observed Rin proviral phenotype could potentially be explained by viral manipulation of Rin to block Imd pathway activity. To test this possibility, we inhibited the activation of Imd by co-depleting Rel2 simultaneously along with depletion of Rin, to determine whether inhibition of the antiviral Imd pathway complements the loss of Rin during the PMI.

The depletion of genes targeted was tested the day of the blood-feeding on unfed mosquitoes. When Rin is depleted the transcript abundance of Rel2 is decreased. When we deplete Rel2, we do not observe a change in the transcript level of Rin (Fig. 3C). Therefore, there is a hierarchical effect which goes in one direction where Rin can regulate Rel2 transcript abundance, but Rel2 does not modify Rin transcript level. First, Rin depletion decreased the number of mosquito abdomens positive for ONNV (Fig. 3A) and diminished the number of ONNV particles in abdomen of dsRin-compared to dsGFP-treated mosquitoes (Fig. 3B). Therefore, Rin depletion decreases the infection rate and the infection intensity of ONNV in An. coluzzii. Rin is required for full levels of both viral infection prevalence and titer at 3 days post-blood feeding in the PMI. Therefore, Rin has a proviral role during the PMI and acts as a switch between a sterile immunity and a viral infection in the in vivo model. Rel2 depletion increases the viral titer but does not modify the infection rate. Therefore, Rel2 is required to limit viral titer, but does not limit infection prevalence. This result indicates that the biological demand to create sterile immunity is more demanding for infection prevalence than for the control of the viral titer. We can hypothesize that the depletion of multiples antiviral pathways will be needed to observe a significant increase of infection prevalence. Depleting Rin and Rel2 simultaneously allows complementation of the Rin phenotype indicating the involvement of Rin in the Imd pathway and its effect on ONNV infection. This data suggests the viral manipulation of Rin to suppress the Imd antiviral pathway and to promote viral infection.

**Figure 3.**
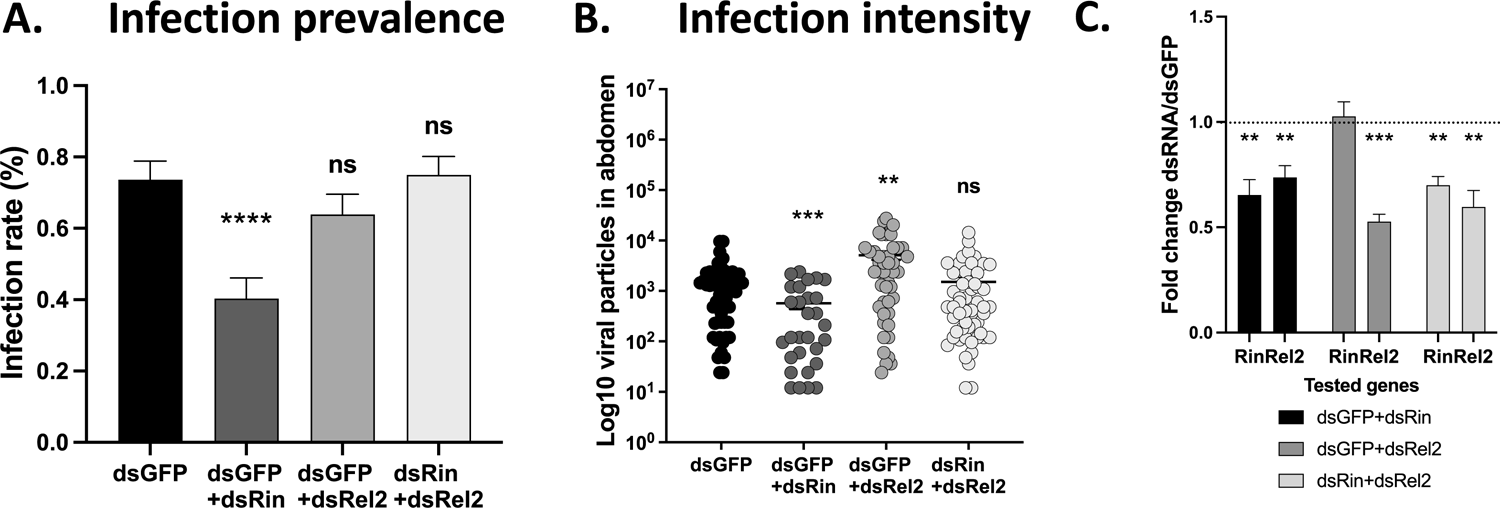
Silencing of the Imd pathway complements loss of Rin during ONNV infection. (A-B). Mosquitoes were treated with dsRNA two days prior to the infectious bloodmeal. Gene silencing was confirmed on the day of infectious blood feeding (C). All fully engorged mosquitoes were sampled at 3 d post-bloodmeal to determine ONNV infection by viral titration of abdomens. (A) ONNV infection prevalence, the proportion of blood-fed mosquitoes positive for infection (B) ONNV infection intensity, the viral titer of only positive mosquitoes. (C) Rin and Rel2 transcript level after dsRNA treatment in unfed female mosquitoes collected the day of the blood-feeding is showed. (A) Each bar represents the percentage of abdomen positive to ONNV between dsGFP, dsGFP+dsRin, dsGFP+dsRel2 and dsRin+dsRel2-treated mosquitoes. (B) Each point represents the viral titer counted in one female mosquito abdomen. (C) Each bar represents the level of transcript abundance of Rin and Rel2 relative to dsGFP-treated control (defined as 1.0). Errors bars indicate the SEM. Each histogram represents the results of 3 independent experiments with 72 mosquitoes analyzed by conditions in total (A-B). A pool of at least 6 mosquitoes has been analyzed by biological replicates (C). ** P < 0.01, *** P < 0.005, **** P < 0.0001, ns non-significant.

### Rin is required for correct basal immunity in Anopheles hemocyte-like cells

After having detailed the role of Rin in the antiviral immunity in the PMI, we know investigate the role of Rin in the systemic compartment using the Anopheles 4a3A hemocyte-like cells. This model has previously been described as a suitable model for the study of the DI (Carissimo *et al*., 2015; Waldock et al., 2012). First, we determined the influence of Rin on ONNV infection at a low multiplicity of infection (MOI: 0.01) in 4a3A cells. Depletion of Rin transcript significantly decreases the intracellular quantity of ONNV genomic RNA and infectious particles (Fig. S1C and S1D). Therefore, Rin activity is proviral for ONNV infection in the DI in the 4a3A cell line.

We analyzed the role of Rin on the transcriptional expression of the same immune genes tested in the in vivo model, in the 4a3A hemocyte-like cells model. We analyzed the transcript fold change of those immune genes between dsRin and dsGFP cells. First, the depletion of Rin was deleterious for the transcript abundance of Rel1, positive regulator of the Toll pathway and APL1C. Rin depletion did not modulate the transcript abundance of TEP3. Thus, Rin increases Rel1 and APL1C transcript expression in 4a3A cells. We also analyzed Rin influence on the Imd pathway (Fig. 4B) and Rin depletion did not influence Rel2 and Rel2F transcript abundance but did reduce APL1A and TEP4 transcript abundance. Rin increases the transcriptional expression of two extracellular immune factors regulated by the Imd pathway, APL1A and TEP4. We then analyzed Rin depletion phenotype on the transcript abundance of other known (LRIM1) or potential immune complex partners subunits (LRIM4, LRIM10 and TEP12) (Fig. 4D). When Rin is depleted LRIM4 and LRIM10 transcript abundance but not LRIM1 and TEP12, are significantly reduced. Thus, Rin enhances the transcriptional expression or the transcript stability of two leucine-rich repeat transcript LRIM4 and LRIM10. The transcript level of the positive regulator of JAK/STAT, Stat-A, is not regulated by Rin (Fig. 4B). Concerning the siRNA pathway, after Rin depletion, Ago2 but not Dcr2 transcript abundance is significantly reduced, thus Rin enhances Ago2 transcriptional expression (Fig. 4C).

**Figure 4.**
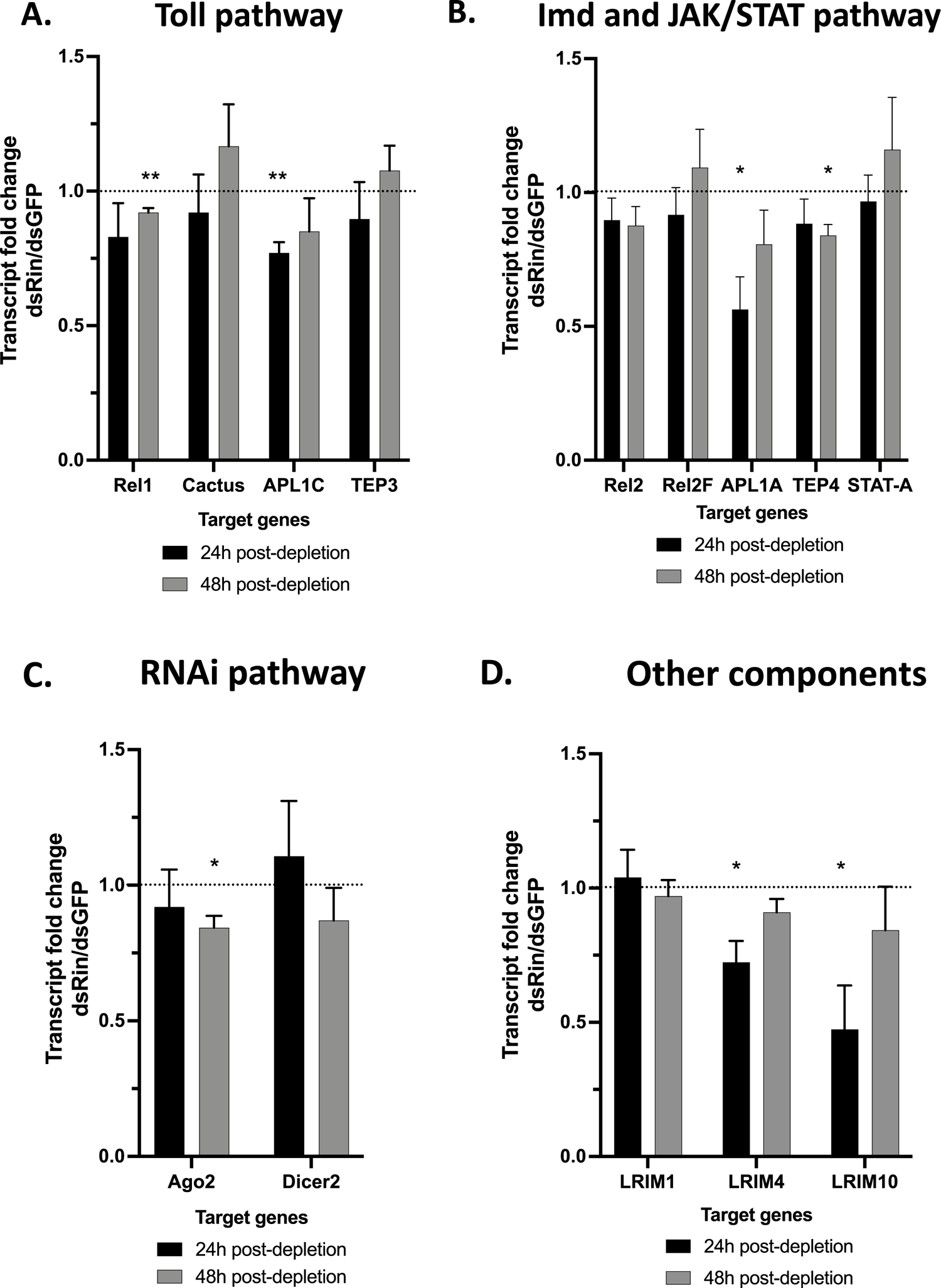
Rin is required for correct basal immunity in Anopheles hemocyte-like cells. (A-D) Cells were depleted for Rin (dsRin) or GFP (dsGFP) using 25 ng/µL of dsRNA per conditions. Rin depletion on genes involved in the Toll pathway (A), the Imd pathway (B), the JAK/STAT pathway (B), the siRNA pathway (C) and other immune transcript involved in complement-like immunity including LRR and TEP transcript (D) are shown. Each bar represents the level of transcript abundance of each immune genes relative to dsGFP-treated control (defined as 1.0), and error bars indicate the SEM. Histogram showed are the results of three independent experiments. * P < 0.05, ** P < 0.01.

These results showed that Rin plays a role in antiviral immunity by positively regulating the transcriptional expression or the transcript stability of multiple immune genes including some previously proven to have antiviral activity. Moreover, Rin modulation of those immune transcript is time-dependent as some were significantly reduced at 24h- or 48h-post depletion indicating respectively an early or a late influence. As some immune transcript abundance came back to the wild-type level at 48h-post depletion we hypothesize that other mechanism may complement the loss of Rin as immune genes are known to be regulated by multiple pathways and partners. Furthermore, Rin mediates transcripts level of immune genes and regulators of NF-kappa B-like pathway in mosquito cells and components of the siRNA pathway.

Moreover, multiple differences are noticed with what was observed in the in vivo model. Indeed, Rin seems to primarily enhance Toll-related immune transcript in the 4a3A systemic compartment model whereas in mosquitoes in vivo model Rin modulates mostly Imd-related immune transcripts. Moreover, other differences observed in the in vivo model compared to the in vitro model can also be linked to the presence of multiple cell types in mosquitoes leading to a higher variability of response than in the one-cell type in vitro model.

### Presence of Rin allows ONNV-dependent inhibition of the Toll pathway in Anopheles hemocyte-like cells

Rin is a proviral factor for ONNV infection in 4a3A cell line (Fig. S1C and S1D) and increases the transcript level of Rel1 the positive regulator of the Toll pathway (Fig. 4A). The Toll pathway in 4a3A systemic model is inhibited by viral infection and among the immune pathways, the Toll pathway artificial stimulation has the most antiviral activity against ONNV replication (Carissimo *et al*., 2015). We investigated the transcript level of the immune genes tested in Fig. 4 during ONNV infection where Rin expression is depleted in 4a3A cells (Fig. 5A). We infected dsRin and dsGFP-treated cells with ONNV at a low multiplicity of infection (MOI: 0.01). Then we analyzed the transcript abundance of those genes between dsRin-treated cells related to dsGFP control-treated cells at 24h- and 48h-post-infection. Of all the genes tested in naïve cells, only Rel1 transcript abundance is modified after Rin depletion in ONNV DI (Fig. 5A). Rin depletion in infected cells increases the transcript abundance of Rel1 at 24h and 48h post-infection compared to infected dsGFP-treated cells (Fig. 5A). Therefore, Rin in infected cells decreases the Rel1 transcript abundance. We wanted then to investigate if the Toll pathway is antiviral in 4a3A cells by depleting Cactus, the negative regulator of the Toll pathway, which activates the Toll pathway (Frolet et al., 2006). We infected dsCactus and dsGFP-treated cells with ONNV at MOI: 0.01. Then we analyzed the intracellular ONNV viral RNA level and the ONNV infectious particle produced in the supernatant between dsCactus-treated cells related to dsGFP-treated cells at 24h and 48h post-infection. Cactus depletion decreases ONNV viral RNA quantity and infectious particles production (Fig. 5B). Therefore, the Toll pathway has an antiviral activity against ONNV infection in 4a3A cells. We previously showed that Rin in uninfected cells is able to upregulate the transcript level of Rel1. However, in infected cells Rin decreases Rel1 transcript level. Consequently, Rin ability to modulate Rel1 expression seems to be manipulated during a viral infection in order to inhibit the antiviral Toll pathway activation in the DI.

**Figure 5.**
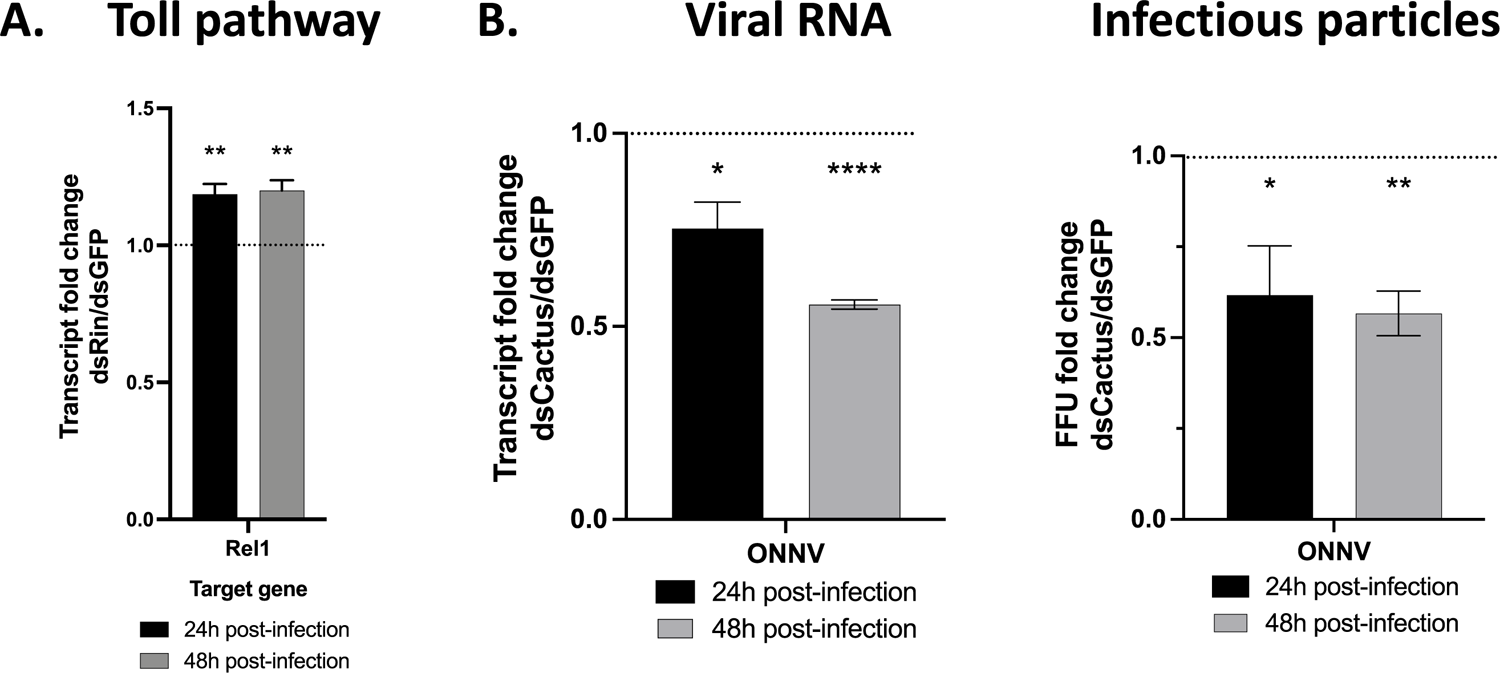
Presence of Rin allows ONNV-dependent inhibition of the Toll pathway in Anopheles hemocyte-like cells. (A-B) 4a3A cells were depleted for Rin (A) or Cactus (B) and infected with ONNV at MOI 0.01. The influence of Rin depletion on Rel1 during a viral infection was tested (A). The antiviral effect of the activation of the Toll pathway via Cactus depletion on ONNV viral RNA quantity and ONNV particle production are presented (B). (A) Each bar represents the level of transcript abundance of Rel1 (A) relative to dsGFP-treated control (defined as 1.0). (B) Each bar represents either the level of transcript abundance of ONNV RNA or focis forming unit relative to dsGFP-treated control (defined as 1.0). Error bars indicate the SEM. Histogram showed are the results of three independent experiments. * P < 0.05, ** P < 0.01, **** P < 0.0001

### Rin physically interacts with ONNV nsP3 in Anopheles hemocyte-like cells

In mammalian and in mosquito cells, G3BPs and Rin have been showed to colocalize with the non-structural protein 3 (nsP3) of multiple alphaviruses including CHIKV and ONNV (Nowee *et al*., 2021). However, it has never been shown in Anopheles cells models. Here we analyzed Rin of An. coluzzii colocalization and interaction with nsP3 of ONNV in 4a3A cells (Fig. 6). We co-transfected a C-terminal streptavidin tag construct of Rin (Rin-strep) and a construct expressing nsP3 of ONNV both controlled by the Anopheles actin promoter (nsp3 transfection). We also directly infected cells with ONNV at MOI 0.5 (ONNV infection). Colocalization pattern was observed between Rin and nsP3 of ONNV in the different conditions. Quantification of the percentage of cells harboring colocalization pattern revealed more than 78% of cells counted showed a colocalization between Rin and nsP3 of ONNV (Fig. 6B). No significant difference was observed between the two conditions.

**Figure 6.**
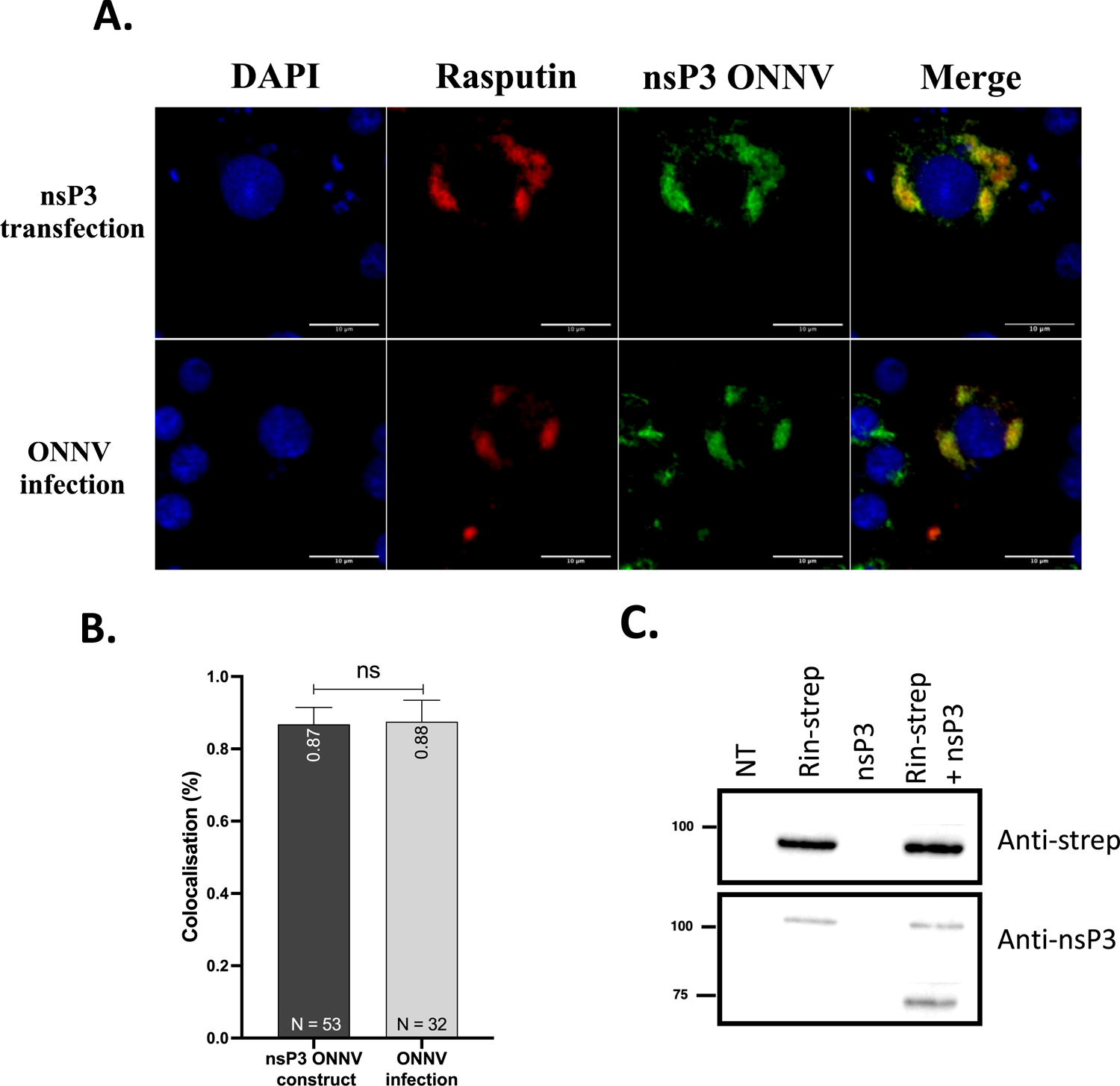
Rin physically interacts with ONNV nsP3 in Anopheles hemocyte-like cells. (A-B) Rin proviral actions on viral RNA quantity (A) and infectious particle production (B) are showed. (C-D) NsP3 signal (green), Rin-strep signal (red), nucleus signal (blue) and colocalization between nsP3 and Rin-strep (yellow) are presented (C). The quantification of the colocalization is showed where the number of cells harboring the colocalization pattern (yellow) are related to the number of co-transfected cells (D). (E) The eluats of the pull-down have been analyzed by Western Blot and bands revealed with antibodies targeting streptavidin tag (anti-strep) and bands revealed with antibodies against nsP3 (anti-nsp3) are represented (E). (A-B) Each bar represents the level of transcript abundance of ONNV RNA (A) or focis forming unit (B) relative to dsGFP-treated control (defined as 1.0), and error bars indicate the SEM. Histogram showed are the results of three independent experiments. *** P < 0.005, **** P < 0.0001. (C-D) A minimum of 30 co-transfected cells were counted by conditions. Representative pictures are showed. Ns non-significant. (E) The result of four independent experiments is showed.

To confirm Rin and nsP3 interaction, we used a streptavidin-pull down assay to purify Rin-strep (Rin-strep) from co-transfected 4a3A cells. We revealed by using streptavidin-specific antibodies and nsP3-specific antibodies, the presence of streptavidin tag protein (Rin) and nsP3 in the eluat of non-transfected cells (NT), Rin-strep transfected cells (Rin-strep), nsP3 transfected cells (nsP3) and co-transfected cells with Rin-strep and nsP3 of ONNV (Rin-strep + nsP3) (Fig. 6C). We observed bands at 89 kDa corresponding to Rin-strep in the eluat of cells transfected with Rin-strep construct or co-transfected with both Rin-strep and nsP3 of ONNV. A band at 70 kDa corresponding to nsP3 of ONNV was only revealed in the eluat of co-transfected cells but was not observed in the eluat of nsP3-transfected cells. Therefore, nsP3 presence in the eluat is due to its binding to Rin-strep (Fig. 6C). Finally, two non-specific bands at 110 kDa are revealed by the nsP3-polyclonal antibodies. Therefore, in 4a3A in vitro model, Rin is a proviral factor for ONNV infection and is also able to physically interact with nsP3 of ONNV similarly to what was known in mammalian cells.

### Rin interaction with ONNV nsP3 modifies the biological function of its partners

Rin presence is required for suppression of the Imd and Toll antiviral pathways during ONNV infection in the PMI and the DI respectively, through a likely virus manipulation of host Rin. Rin interacts directly with the nsP3 of ONNV in cells, therefore this interaction could be at the center of the coopting of Rin. To shed light on the mechanism by which Rin regulates antiviral genes, we investigated the protein partners of Rin and of the complex Rin/nsP3 in 4a3A cells. We analyzed by mass spectrometry the elution of cells transfected with Rin tagged with a C-terminal streptavidin sequence (Rasputin-strep), or the elution of co-transfected cells with Rin-strep and nsP3 of ONNV (Rasputin-strep+nsP3) (Fig. 7). Controls were used to attest of the specificity of the partners found. Among these controls, the elution of cells transfected with GFP-strep or transfected with nsP3 or non-transfected was also analyzed by mass spectrometry. Proteins found in the control conditions are considered non-specific to Rin-strep and are, therefore, removed from the data set.

**Figure 7.**
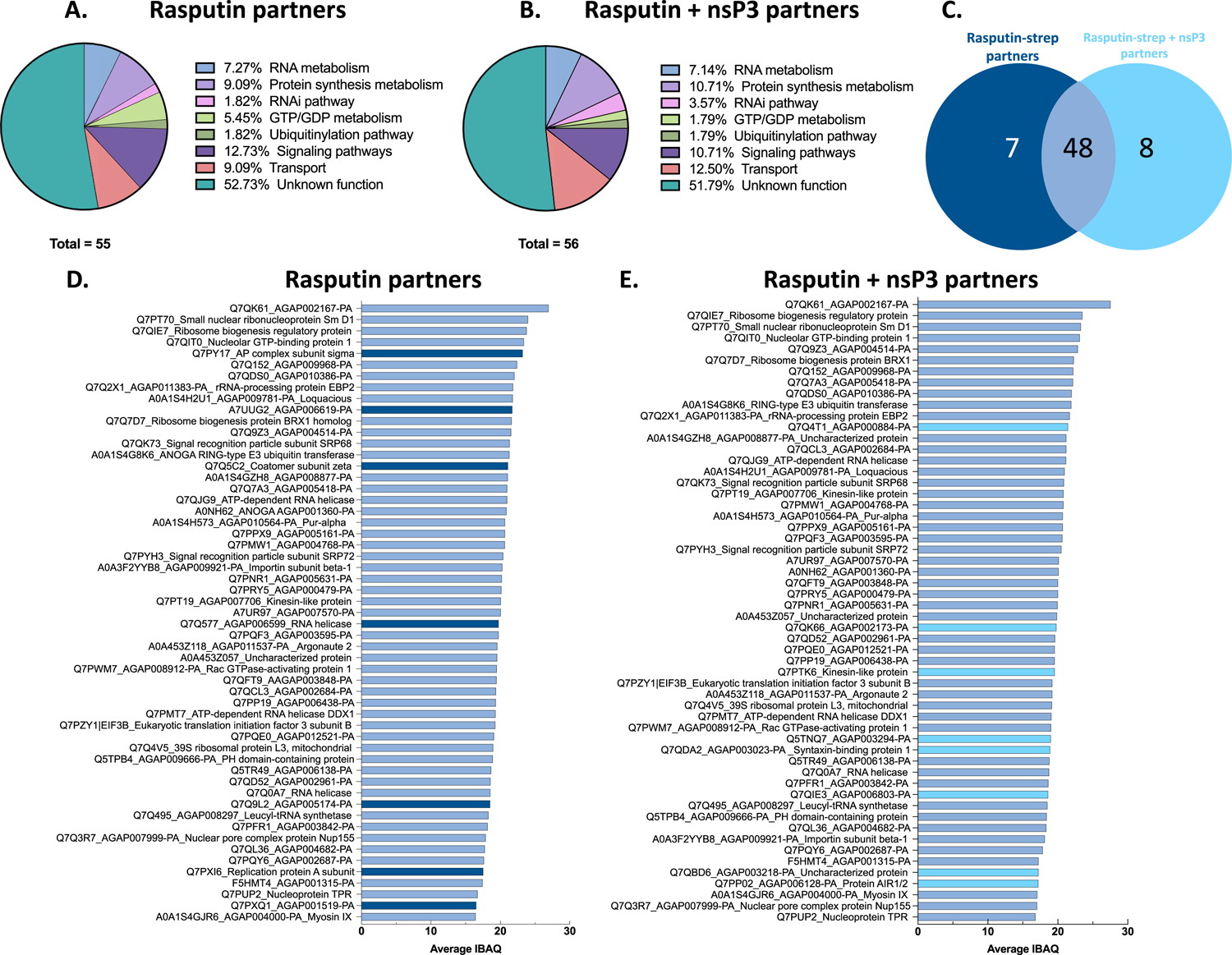
ONNV nsP3 binding alters Rin protein partners in Anopheles cells. (A-E) Eluats of cells transfected with Rin-strep or co-transfected with Rin-strep and nsP3 of ONNV were analyzed by mass spectrometry to identify proteins partners. The non-specific proteins found in the controls conditions were removed from the results. (A) Summary of the biological functions of the protein partners of Rin-strep, or (B) of Rin-strep in presence of nsP3 are showed. (C) A summary of the partners proteins in common between Rin-strep and the complex Rin-strep/nsP3 is presented. (D-E) The average IBAQs of the protein partners of Rin-strep (dark blue) or Rin-strep/nsP3 complex (light blue) or their protein partners in common (plain blue) are showed. Each bar represents the average IBAQs of each proteins partners of Rin-strep or of the complex Rin-strep/nsP3 where the non-specific proteins have been previously removed. Data showed are the results of four independent experiments per conditions.

Most Rin partners are proteins with unknown function. Of the known proteins, Rin interacts in diverse pathways including RNA metabolism, protein synthesis components, ubiquitination pathways and signaling pathways (Fig. 7A). The biological pathways of Rin partners are in line with Rin involvement in Ras metabolism (Pazman et al., 2000) and RNA metabolism (Laver *et al*., 2020). When Rin is in complex with the viral nsP3, more proteins partners of Rin are involved in protein synthesis, RNAi pathway and cellular/nuclear transport (Fig. 7B). However, the interaction between Rin and nsP3 decreases the number of Rin partners involved in GDP/GTP metabolism and in signaling pathways. Therefore, the presence of nsP3 modifies the biological profile of Rin protein partners. Interestingly, Rin does not directly interact with the immune genes that it regulates except Ago2. Consequently, Rin modulation of the transcript levels of antiviral genes most likely derives from Rin function in RNA metabolism or protein synthesis as it has been already reported (Laver *et al*., 2020). Moreover, 48 proteins were found to interact with Rin in the presence or absence of nsP3 whereas 7 proteins interact only with Rin alone and 8 proteins interact only with Rin when nsP3 is present (Fig. 7C). Therefore, the presence of the viral nsP3 allows Rin to still interact with most of its protein partners but also modifies the partner profile of Rin. Consequently, the interaction between Rin and the viral nsP3 could manipulate the protein partners of Rin and therefore may influence Rin function in modulating antiviral genes.

## DISCUSSION

The mechanism driving the proviral role of G3BPs on alphavirus infection has primarily been studied during the viral RNA replication steps of the viral genome (Gotte et al., 2019; Gotte *et al*., 2020; Scholte *et al*., 2015). G3BPs in mammals have been shown to influence the switch from genome replication to genome translation (Scholte *et al*., 2015). However, as we know that G3BPs have multiple functions, other potential roles of G3BPs during the alphavirus cycle have not been investigated yet. Moreover, the role of G3BPs in immune pathways has already been mentioned especially in NF-kappa B signaling pathways. But the relationship between G3BPs and immunity during a viral infection has never been made. Therefore, this study reveals a link between Rin, immunity and viral infection and could be extrapolated to Rin in other mosquito species and to G3BPs in mammals.

Rin positively regulates immune genes and known antiviral genes with a centered effect on the Toll pathway in the systemic compartment represented by the 4a3A cell line (Fig. 4) and a pronounced effect on the Imd pathway in the midgut compartment of the in vivo model (Fig. 1). These results are in adequation with what was observed in Carissimo et al, regarding the immune compartmentalization of these pathways during ONNV infection. Where the Toll pathway is mostly antiviral in the DI whereas the Imd pathways is mostly antiviral during the PMI (Carissimo *et al*., 2015). Moreover, Rin transcriptional expression is enhanced following a naïve blood-feeding (Fig. S5) which is in line with our observation of Rin enhancing multiple immune factors. Indeed, during bloodfeeding, mosquitoes will be exposed to the pathogens found in the blood of the mammals fed upon. Thus, mosquitoes will need to restrict the infection of these pathogens for their survival by enhancing the activation of these immune pathways.

Surprisingly, Rin effect is not consistent in time. Most of the genes regulated by Rin are either lately modulated or come back to wildtype level later after Rin depletion (Fig. 1 and 4). These results indicate the plurality of the regulation of those immune genes and the possibility that other proteins could intervene in the Rin-mediated immune gene regulation. Indeed, most of those genes are regulated by multiple factors and the effectors of some immune pathways could be regulated by other immune pathways. Moreover, differences of immune gene response following Rin depletion observed between the in vitro and in vivo models can be explained by the plurality of the cell types present in the mosquito model compared to the one single cell type of the in vitro 4a3A cell model (Muller et al., 1999). Indeed, G3BPs are known to be ubiquitously expressed in multiple human tissues. As for Rin in mosquitoes no study examined the expression of Rin in different tissues of mosquitoes (Kennedy et al., 2001), but the mammalian studies could be extrapolated to mosquitoes as G3BPs/Rin are overall well-conserved proteins (Kang *et al*., 2021). Therefore, Rin could mediate various genes depending on the cell-type where it is expressed.

Interestingly, we found that during the PMI, multiple immune factors were upregulated when Rin was absent compared to a wt viral infection (Fig. 2). Taking in account that in mosquitoes Rin allows the switch between sterile immunity and infection during the PMI, central point for the establishment of a viral infection (Fig. 3), these results demonstrate a correlation between the proviral role of Rin during viral infection and Rin immune pathway regulation. Indeed, in non-infected mosquitoes Rin can upregulate the expression of immune genes but during a viral infection Rin decreases the expression of those same immune genes (Fig. 1). Therefore, Rin physiological function in naïve mosquitoes is corrupted by the viral infection. Furthermore, the linked between Rin immune gene regulation and viral infection was directly made in mosquitoes via the complementation of Rin deleterious phenotype on the viral infection by the inhibition of the Imd pathway (Fig. 3). This result directly proves that Rin proviral function is directly linked to its ability to regulates the Imd pathways. Rin activity is required to overcome a wildtype cellular qualitative (prevalence) and quantitative (titer) blockade against ONNV infection (Fig. 3). The depletion of Rel2 complements the loss of Rin activity by reversing the effect of the cellular blockade and restoring normal ONNV infection prevalence and titer levels. This result suggests that Imd pathway activity comprises and explains at least in large part the PMI blockade and is consistent with a mechanism in which ONNV interacts with Rin to subvert Imd activity and promote viral infection. In the absence of Rin activity, the antiviral phenotype of Rel2 is enhanced, decreasing ONNV infection, while in the absence of Rel2, the proviral immune subversion function by Rin is not required for normal levels of ONNV infection. Moreover, we revealed that there is a hierarchical relation between Rin and Rel2 where Rin is able to regulate the transcript level of Rel2, but Rel2 cannot regulate Rin transcript abundance back (Fig. 3). This observation supports the evidence of a directional effect mediates by Rin on this antiviral pathway where Rel2 is downstream of Rin in this chain of event.

The simplest model for the relationship between infection prevalence and viral titer could be that the activity of predominantly the Imd pathway reduces the efficiency of viral replication below a level that can consistently yield an ongoing infection, and sterile immunity results in those individuals where a combination of low viral titer and the action of other antiviral and stochastic effects allows elimination and curing of the viral infection. The depletion of Rel2 alone allowed a significantly elevated viral titer but did not alter infection prevalence. It is not clear why the silencing of only Rel2 did not alter ONNV infection prevalence, while the silencing of Rel2 in the absence of Rin complemented the loss of Rin by reversing the depleted Rin phenotype. The most likely explanation is that infection prevalence in the dsGFP-treated control mosquitoes was close to 80%, and thus the experimental design may have been powered to detect decreased prevalence but lacked power to detect a significant increase above 80%. An alternative explanation could be that depletion of different antiviral pathways such as Imd or JAK/STAT can individually result in elevated viral titer, but that complete elimination and sterile immunity requires the integrated effect of multiple antiviral pathways. Further work would be required to distinguish between these possibilities. Taken together, the results indicate that the proviral effect of Rin for ONNV infection is mediated at least in large part by a Rin-dependent inhibition of Imd pathway activity.

Other viral mechanisms could be involved in the stabilization of the expression of those immune genes. Moreover, Carissimo et al highlighted a possible inhibitory activity of ONNV against the Toll pathway in 4a3A cells (Carissimo *et al*., 2015). Here we showed that the Toll pathway stimulation decreases the viral infection therefore, Rin regulation of this pathway make it a target of choice for ONNV. This study may suggest that Rin would be at the center of this viral inhibitory of the Toll pathway during the DI. However, extensive study will be required to directly prove the Rin proviral function role via the Toll pathway regulation. Directly studying the role of Rin on the DI in the in vivo model would be possible via the intrathoracic injection of viral particles in Rin-depleted mosquitoes allowing the bypass of the midgut barrier.

We revealed that similarly as what was observed with Rin in Aedes and G3BPs in mammals (Nowee *et al*., 2021), Rin of Anopheles is also able to interact with nsP3 of ONNV (Fig. 6). This result indicates the conservation of this interaction even in the poor arbovirus vector as Anopheles. Moreover, Rin is also clearly proviral for ONNV infection in Anopheles (Fig. S1) and is upregulated during the DI (Fig. S2). Therefore, Rin/G3BPs and alphavirus interaction is overall conserved between invertebrates and vertebrates, which highlight the importance of this relationship for the viral cycle. Viruses have evolved to strongly interact with Rin/G3BPs both in invertebrate and vertebrate hosts. Therefore, this interaction between nsP3 and Rin/G3BPs is strongly needed for the virus to be maintained in both hosts. This evidence strongly suggests an important role for Rin during the viral cycle. As previously mentioned, it would be surprising that Rin role, knowing its plurality of functions, is only used by viruses during their RNA replication steps. Our hypothesis is that ONNV nsP3 will hijack Rin for both replication step functions and for inhibiting the innate antiviral immunity of its host.

The study of Rin partners revealed numerous protein partners involved in various biological function mostly RNA metabolism and protein synthesis. However, the siRNA component Ago2 is also a partner of Rin. Moreover, most of Rin partners have no known functions which highlighted the complexity of the Rin network. Furthermore, the presence of nsP3 changes the proportion of Rin partners and the biological function of these partners. These results revealed that the nsP3 interaction with Rin is able to modify the scheme of Rin partners. However, contrary to what was observed in Prigent et al with G3BPs capable to interact with IkB alpha and NF-kappa B/I-kappa B alpha complex (Prigent *et al*., 2000). In the mosquito vector no immune factors except Ago2 was identify as a partner of Rin. Moreover, similarly as our study, Scholte et al mentioned that they did not discover NF-kappa B and I-kappa B alpha transcription factor in their interaction assay with G3BP2 in mammals cells (Scholte *et al*., 2015) Detailed studies need to be done to specifically dissect the molecular mechanism by which Rin influence the NF-kappa B antiviral immune pathways Toll and Imd. Our hypothesis is that Rin would allow the regulation of these immune genes by intervening in the transcription, RNA stabilization, or protein synthesis of regulatory factors in those antiviral pathways. To confirm this hypothesis, an interaction assay between Rin and mRNA in cells will be necessary in order to identify the mRNA partners of Rin.

This study revealed the link between Rin, antiviral immunity and viral infection and could be extrapolated to the role of G3BPs in vertebrate cells. Although Rin was always mostly studied for their role in the RNA replication steps of the virus, even if no specific mechanism was explained yet, the plurality of G3BPs functions in the cell metabolism make them a target of choice for the virus to manipulate. Here we highlighted that ONNV hijacks Rin probably via its interaction with nsP3 in order to corrupt the antiviral immune pathway that Rin regulates. Although Rin is a choice target for the virus, its plurality in multiple biological pathways would make it a difficult target for therapeutic treatment with the risk of interfering with other function in the organism. However, dissecting the partners of Rin with unspecified function could point to promising new therapeutic targets.

## MATERIALS AND METHODS

### Cellular strains, viral strains, and mosquitoes

4a3A cells originated from neonate An. coluzzii 4a r/r strain larvae spontaneously immortalized (Muller *et al*., 1999). 4a3A cells were maintained at 27°C without CO_2_ in Insect-XPRESS™ Protein-free Insect Cell Medium complemented with heat-inactivated Fetal Calf Serum (FCS) (#A3840002 ThermoFisher Scientific). C6/36 cell line (ATCC) (Igarashi, 1978) was used for viral titration and were maintained in Leibovitz’s L-15 medium (#11415-049, ThermoFisher Scientific) supplemented with MEM non-essential amino acid (#11140-050, ThermoFisher Scientific) and complemented with heat-inactivated Fetal Calf Serum (#A3840002 ThermoFisher Scientific) at 28°C without CO_2_. Baby hamster kidney cell line (BHK-21) was used to produce viral particles from infectious clones and were maintained in Dulbecco’s Modified Eagle Medium (DMEM) (1X) + GlutaMax media (#61965-026, ThermoFisher Scientific) complemented with heat-inactivated Fetal Calf Serum (#A3840002 ThermoFisher Scientific) at 37°C with 5% CO_2_.

Infectious clone of ONNV used in this project originated from ONNV strain Igbo Ora-IBH10964 (Igbo-Ora strain) and was isolated from a human febrile patient during the epidemic of 1966 in Nigeria (accession number AF079457) (Lanciotti et al., 1998). ONNV infectious clone was either produced on Anopheles 4a3A cells and viral titer done on kidney epithelial cells from African green monkey (Vero cells) were measured at 7,3 x 10^5^ pfu/mL (plaque forming unit) for cellular infection. Mosquito infection used infectious ONNV clone produced on BHK-21 cells and the titer measured on C6/36 cells is 5,64 x 10^7^ ffu/mL. Infectious clone cDNA was linearized by PmeI, and viral RNA was transcribed using T7 RNA polymerase (Ambion mMESSAGE mMACHINE® SP6 Kit). Transcribed RNA was electroporated into BHK-21 or 4a3A cells. Virus was recovered after 72 h and titer on C6/36 cells.

The An. coluzzii Ngousso colony was created from 100 wild-caught female mosquitoes from Cameroon in 2006 and are bred at the Institut Pasteur Center for the Production and Infection of Anopheles (CEPIA) facility since 2008. All the steps of mosquitoes breeding occurred in a controlled environment with a temperature of 26°C (±1°C), 70% humidity and 12h:12h of day: night photoperiod. Female mosquitoes used for all experiments are of the second generation (F2) and are of early emergence (less than 5 days post-emergence).

### Primers and plasmids

The dsRNA sequences were either previously published or constructed using the cDNA sequence of genes in the An. gambiae PEST genome available on VectorBase database. DsRNAs targeting GFP act as a control for the dsRNA treatment. DsRNA were synthesized using T7 primers and the transcription kit of MEGAscript™ T7 (#AM1334, Invitrogen) following manufacturer instructions. DsRNA were used either in 4a3A cell line or in An. coluzzii Ngousso mosquitoes.

Plasmids used for colocalization, and streptavidin pull-down assay were constructed using Rin codon-optimized AGAP000403 sequences and ONNV nsP3 protein from IBH10964 strains graciously given to us by Andres Merits. A C-terminal streptavidin tag II (W-S-H-P-Q-F-E-K) was added at the end of Rin sequences. For their expression in 4a3A cells, these genes are expressed under the Actin promoter (Ac5) in the plasmid pAc5.1. 4a3a cells lines were transfected using lipofectamine LTX (#A12621, ThermoFisher Scientific) with 0,25 µg to 0,5 of plasmids depending on the number of cells used.

### RNA extraction, cDNA synthesis and qPCR analysis

Following experiments, RNA was extracted from 4a3A cells or mosquitoes using Trizol reagent (#T9424, Sigma Aldrich) and RNA miniprep kit (#ZR1054, Ozyme). Complementary DNAs were then synthesized with M-MLV transcriptase inverse kit (#28025013, Invitrogen) using 1 µg of RNA. The qPCR primers were checked for specificity prior to the analysis. All qPCRs were performed using SYBR green supermix (KAPA SYBR FAST ABI, Sigma-Aldrich) and the CFX96 Touch Real-Time PCR Detection System (Biorad). The ribosomal protein rpS7 gene was used as the internal control and the analysis of transcript relative expression to rpS7 was performed according to the 2−ΔΔCt method. Conditions of the run were: 95°C for 10min, then 39 cycles of 95°C for 15 sec, 60°C for 1min (plateread).

### Antibodies and fluorescent labels

Sera of rabbit containing polyclonal rabbit antibodies graciously given to us by Andres merits targeting the macro and the AUD domain of nsP3 of ONNV were used at 1:100 overnight at 4°C for colocalization and western blot assay. Secondary Goat anti-Rabbit IgG (H+L) antibodies (#A27039, Invitrogen) conjugated with Alexa Fluor 555 were used during colocalization assay at 1:500 for 45 min at room temperature (RT) 0,2% Triton X-100 1% BSA PBS 1X. Secondary antibodies anti-rabbit conjugated with HRP (#7074S, Invitrogen) were used for western blot at 1:8000 for 1 hour at RT in 5% BSA in TBS1X solution. Monoclonal antibodies targeting the streptavidin tag II and conjugated with HRP (#2-1502-001, IBA) were used at 1:10000 for 1h at RT during western blot in 5% BSA in TBS1X solution. Streptavidin-tactin molecule conjugated with dye-649 (#2-1568-050, IBA) were used at 1:100 for 3h at RT during colocalization assay in 0,2% Triton X-100 1% BSA PBS 1X. DAPI (4’, 6-diamidino-2’-phenylindole, dihydrochloride) (#62247, ThermoScientific) was used at 1:1000 for 10 min at RT in PBS1X for staining the nucleus of 4a3A cells during colocalization assay. Revelation of viral titration was made using monoclonal mouse antibodies against CHIKV Virus-like particles (#MAB12385, The Native Antigen), diluted at 1:1000 in 0,1% BSA PBS1X solution for 45 min at 37°C. Secondary goat antibodies against mouse IgG and IgM (H+L) conjugated with Alexa 488 (#A-10680, Invitrogen) were diluted at 1:500 in PBS1X and incubated at 37°C for 30 min.

### Gene silencing

For silencing gene in the *in vivo* system, 500 ng of dsRNA in a final volume of 70 nL maximum was injected into the thorax of cold-anesthetized 1-to 2-d-old An. coluzzii females using Nanoject II injector (Drummond Scientific) and glass capillary needle as previously described (Mitri *et al*., 2009). The gene silencing efficiency was verified after dsRNA injection with RNA from a pool of six un-fed whole mosquitoes by RT-quantitative PCR (qPCR). For silencing genes in the *in vitro* system, a total of 1 × 105 An. coluzzii 4A3A cells were seeded and left to adhere overnight. Cells were incubated for 30 min on a rocker with 500 ng of in 200 µL of Insect Xpress media without FBS. After incubation, 300 µL of microliters of Insect Xpress media with 10% (vol/vol) FBS was added, and cells were incubated until collection or infection. Gene silencing efficiencies were confirmed and are shown in Fig S3 and Fig S4.

### Cellular infection

The 4a3A cell line were infected with ONNV at multiplicity of infection (MOI) of 0.01 (for dsRNA experiment) or at MOI 0.5 (for colocalization experiment) for 1h at 28°C without CO2 in Insect-X-Press media without FCS. After 1 hour of infection, cells were washed three times with Insect-X-Press without FCS. Then 2%FCS Insect-X-Press media was added to cells for the time of incubation. Cells were then collected at 24h or 48h post-infection.

### Infectious blood feeding

The experimental protocol is based on published methods (Vazeille et al., 2007). Infectious blood meal was composed of 2/3 rabbit erythrocytes and 1/3 of viral solution for a final titer of 10^7^ ffu/mL. Adenosine triphosphate (ATP) at 10mM was also added to the infectious blood meal. The infectious blood meal has been distributed in capsule cover with pork intestine. Blood-filled capsules were heat at 37°C on Hemotek® system. Mosquitoes were feed on infectious blood meal for 1 hour. After that female mosquitoes were anesthetized on ice and blood-stuff females were incubated at 28°C and 80-90% humidity with water-imbibed cotton with 10% sugar. Abdomens and thoraces of female mosquitoes were separated, grinded at 2000 rpm for 30 seconds and centrifuged at 4°C for 5min at 10 000 rpm. Supernatants was collected and conserved at −80°C until analysis.

### Viral titration by focus forming unit

Ae. albopictus C6/36 cells were seeded in 96-well plates at 1.25 x 10^6^ cells/ Leibovitz’s L-15 medium (#11415-049, ThermoFisher Scientific) supplemented with MEM non-essential amino acid (#11140-050, ThermoFisher Scientific) with 10% of FBS and incubated at 28°C without CO_2_, 48h prior to the titration. Virus-containing samples were sequentially diluted and 50µL of those dilutions were added to cells in one well. After one hour of incubation at 28°C, 150 µL of 1:1 mixed with 4% carboxymethylcellulose (CMC) with L-15 containing 5 % of FBS is added on cells. Antibiotic-antifungal solution (#15240062, Life technologies®) is also added at a finale concentration of 1.5X. Plates are incubated at 28°C without CO_2_ for 3 days, then cells are fixed using 3.4% formaldehyde in PBS1X for 20 min at room temperature. Cells are then washed three times in PBS1X and then stained with primary antibodies. After three washes with PBS1X, cells are stained with secondary antibodies. Viral titers are counted by observing foci using a fluorescent microscope and the virus titer of the tested sample is expressed as foci forming unit/mL (FFU/mL).

### Confocal microscopy

Cells were seeded in µ-slide 8-well plate (#80807, Ibidi) and transfected using lipofectamine with 0.25 µg of plasmids. Two days post transfection cells were fixated with 4% formaldehyde for 20min at RT. Cells were then blocked and permeabilized using a solution of 0,2% Triton X-100 1% BSA PBS 1X for 1h at RT. Then cells were stained with different antibodies at specific dilution in 0,2% Triton X-100 1% BSA PBS 1X. Washes were realized using 0,2% Triton X-100 1% BSA PBS 1X. Colocalization analysis in 4a3A cells were done using a laser-scanning confocal microscope (LSM700, Carl Zeiss Jena) at the UTechS Photonic BioImaging - C2RT platform (Pasteur Institute). All acquisition parameters were done as listed below: 63X oil-objective, frame size 512×512, pixel dwell 1,57 µsec, scan time 5,78 sec, 16-bit depth, image size 32,9×32,9 µm and 0,06 µm pixel size. The DAPI laser (450 nm) had an intensity of 2, a pinhole diameter of 0,6 µm, a master gain of 730 and a digital gain of 1. The Red A555 laser (514 nm) had an intensity of 2,6, a pinhole diameter of 0,6 µm, a master gain of 711 and a digital gain of 1. The Far-Red D647 laser (633nm) had an intensity of 2, a pinhole diameter of 0,8 µm, a master gain of 700 and a digital gain of 1. Images were analyzed using Fiji software (Schindelin et al., 2012). Background signals were calculated with the mean of signals from 5 pictures of the negative controls of each fluorophore. Mean of background signals were then removed from the analyzed pictures. Colocalization was quantified by counting the number of cells with colocalization pattern related to cells co-transfected (harboring both signals). Between 30 and 50 cells were counted in the analysis.

### Streptavidin-pull down assay

First, 0.5 µg of plasmid constructs were transfected in 4a3A cells using lipofectamine LTX with Plus Reagent (#15338030, ThermoFisher Scientific). Two days post-transfection, cells lysis has been performed using cell lysis buffer (containing Tris 1 M, NaCl 5M, EDTA, Igepal, Protease inhibitor and Phosphatase inhibitor) for 30 min at 4°C and samples were centrifuged at 16,000g for 15 minutes at 4°C. Pull-down assay was performed using MagStrep” type”3 beads (Strep-Tactin® XT coated magnetic beads, 5 % (v/v) suspension, # 2-4090-002, IBA) with a binding capacity up to 0.85 nmol/μl beads (corresponding to 25,5 μg of a 30 kDa protein). Beads are first wash with cell lysis buffer then cell lysate is added on the beads and incubated for 2h at 4°C on a tube rotator. After 2 hours of incubation, lysates are removed, and washes are performed using 1X W buffer (#2-1003-100, IBA). Following washes, either on-beads digestion at the Proteomic platform is done for Mass spectrometry analysis or the elution of proteins is performed using 1X BXT buffer containing biotin (#2-1042-025, IBA) for 10 min. Eluates are then analyzed by Western Blot in order to detect strep-tag proteins (Rin or GFP, bait) and partners (nsP3, prey) using specific antibodies.

Proteins samples were reduced in 1X DTT and heat at 95°C for 5min. Then samples are mixed with XT sample buffer (#1610791, Bio-Rad) and loaded on 4-12% Criterion SDS-PAGE gels (#3450124 Bio-Rad). SDS-PAGE is run at 100V for 10 minutes (stacking gel) and 150V for 1h00min (running gel) in NOVEX NUPAGE MOPS SDS Running Buffer 20X (#NP0001, Life Technologies). Following electrophoresis, proteins contained on gel are transferred onto a 0,2 µm nitrocellulose membrane (#1704159, Bio-Rad) using Bio-Rad Trans-Blot® Turbo™ Transfer System. Transfer program is run for 7min at 2,5A and 25V. After transfer, immunoblots are blocked in a solution containing 5% of Bovine Serum Albumin (BSA) called TBS1X (Tris-buffered saline, 0.1% Tween 20). Then immunoblots are probed with different antibodies at specific dilutions in 5%BSA PBS1X. Detection step was performed using the Enhanced chemiluminescence (ECL) system (Clarity Western ECL substrate #170-5060, Biorad) following manufacturer instruction. Blots were photographed and revealed taking one picture every second for 10 seconds using ChemiDoc Imaging System (Bio-Rad). Blots were then annotated, and molecular weights were measured related to the proteins ladder (Precision PlusProtein All Blue Standards #1610373, Bio-Rad) using the Image Lab analysis software (Bio-Rad). A positive control is performed using a commercial GFP-strep (Green Fluorescent Protein) (#2-1006-005, IBA-LifeSciences) in order to confirm the success of the Western Blot.

### Mass spectrometry analysis

Following streptavidin-pull down assay, on-bead digestion of sample is performed at the Proteomic platform (CNRS – UAR2024) at Pasteur Institute. Proteins were resuspended in 50mM ammonium bicarbonate pH 8.0 and reduced for 30 min at room temperature (RT) with 5 mM TCEP and then alkylated at 50 mM iodoacetamide (I114 - Sigma, St Louis, Missouri, USA) for 30 min, RT in the dark. Protein was then digested with 0.5 µg Sequencing Grade Modified Trypsin (V5111 - Promega, Madison, Wisconsin, USA). Following the overnight digestion at 37°C, 800rpm in the thermomixer C (Eppendorf), the samples were placed on a magnet for 1 min and the supernatant containing peptides transferred in a new tube. Resulting peptides were acidified at 1% with formic acid, desalted with the AssayMAP Bravo robot using C18 column and AssayMAP Peptide Cleanup Protocol (Agilent). All samples were dried in a Speed-Vac and peptides were resuspended in 2 % ACN, 0.1 % FA prior to LC-MS/MS analysis. LC-MS/MS analysis of digested peptides was performed on an Orbitrap Eclipse mass spectrometer (Thermo Fisher Scientific, Bremen) coupled to an EASY-nLC 1000 (Thermo Fisher Scientific). Peptides were loaded (at constant pressure of 900 bars) and separated at 250 nl.min-1 on a home-made C18, 30 cm capillary column picotip silica emitter tip (75 μm diameter filled with 1.9 μm Reprosil-Pur Basic C18-HD resin, (Dr. Maisch GmbH, Ammerbuch-Entringen, Germany)) equilibrated in solvent A (2 % ACN, 0.1 % FA). Peptides were eluted using a gradient of solvent B (80 % ACN, 0.1 % FA) from 2 % to 7 % in 3 min, 7% to 31 % in 42 min, 31 % to 62 % in 10 min, 62 % to 95 % in 5 min (total length of the chromatographic run was 70 min). Column was equilibrated with 10µl of A at 900 bar. Mass spectra were acquired in data-dependent acquisition mode with the XCalibur xxx software (Thermo Fisher Scientific, Bremen). MS spectra were acquired at orbitrap resolution of 60k (at m/z 400), AGC target 800000, a custom max injection time. The scan range was limited from 300 to 1500 m/z. Peptide fragmentation was performed using higher-energy collision dissociation (HCD) with the energy set at 27 NCE. The MS/MS spectra acquired at a resolution of 30k (at m/z 400). Isolation window was set at 1.6 m/z. All acquisitions were done in profile and positive mode. Dynamic exclusion was employed within 30s.

Proteomic platform has performed data and statistical analysis. The guideline used is described below. Raw data were analyzed using MaxQuant software version 2.0.3.0 (Tyanova et al., 2016) using the Andromeda search engine (Cox et al., 2011). The MS/MS spectra were searched against a custom database from Vectorbase repositories 20PEST database (download in 25/04/2022) and tagged proteins. Usual known mass spectrometry contaminants and reversed sequences of all entries were included. Andromeda search was performed choosing trypsin as specific enzyme with a maximum number of two missed cleavages. Possible modifications included carbamidomethylation (Cys, fixed), oxidation (Met, variable), Nter acetylation (variable). The mass tolerance in MS was set to 20 ppm for the first search then 4.5 ppm for the main search and 20 ppm for the MS/MS. Maximum peptide charge was set to seven and seven amino acids were required as minimum peptide length. The “match between runs” feature was applied for samples having the same experimental condition with a maximal retention time window of 0.7 minute. One unique peptide to the protein group was required for the protein identification. A false discovery rate (FDR) cutoff of 1 % was applied at the peptide and protein levels. For the differential analyses of one condition versus another, proteins identified in the reverse and contaminant databases and proteins “only identified by site” were first discarded from the list of identified proteins. Then, only proteins with at least three quantified intensities in a condition were kept. The proteins of interest are therefore those which emerge from this statistical analysis supplemented by those which are considered to be present from one condition and absent in another.

### Statistical analysis

All experiments presented are the results of at least three independent experiments. Bar plots were created using Prism. Statistical significance of most of the experiments presented was established using a two-tailed unpaired (independent) Student t-test which compare the mean of two independent groups (control and treatment). A two-tailed non-parametric unpaired Mann Whitney test was performed for assessing the statistical significance of the difference of viral titer in mosquitoes. The P values were assessed with a null distribution. The P values were considered significant if p<0.05 (*p<0.05; **p<0.01; ***p<0.005, ****p<0.0001).

### Availability of data and material

All data from the study are presented within this article.

### Ethics statement

The protocol for the ethical treatment of the animals used in this study was approved by the research animal ethics committee of the Institut Pasteur, “C2EA-89 CETEA Institut Pasteur” as protocol number 202195.02. The Institut Pasteur ethics committee is authorized by the French Ministry of Higher Education and Research (MESR) under French law N° 2001-486, which is aligned with Directive 2010/63/EU of the European Commission on the protection of animals used for scientific purposes. The study was performed using practices and conditions approved by the Institut Pasteur Biosafety Committee as protocol number CHSCT 14.114.

## Author contributions

Conceived and designed the research: SC CM ABF KDV. Performed the experiments: SC AB CM EBF MM. Analyzed the data: SC CM MM KDV. Wrote the paper: SC CM ABF KDV.

## Acknowledgments

We thank the Institut Pasteur Center for the Production and Infection of Anopheles (CEPIA) for provision of mosquitoes.

## SUPPORTING INFORMATION CAPTIONS

### Supporting Figures

**Figure S1.**
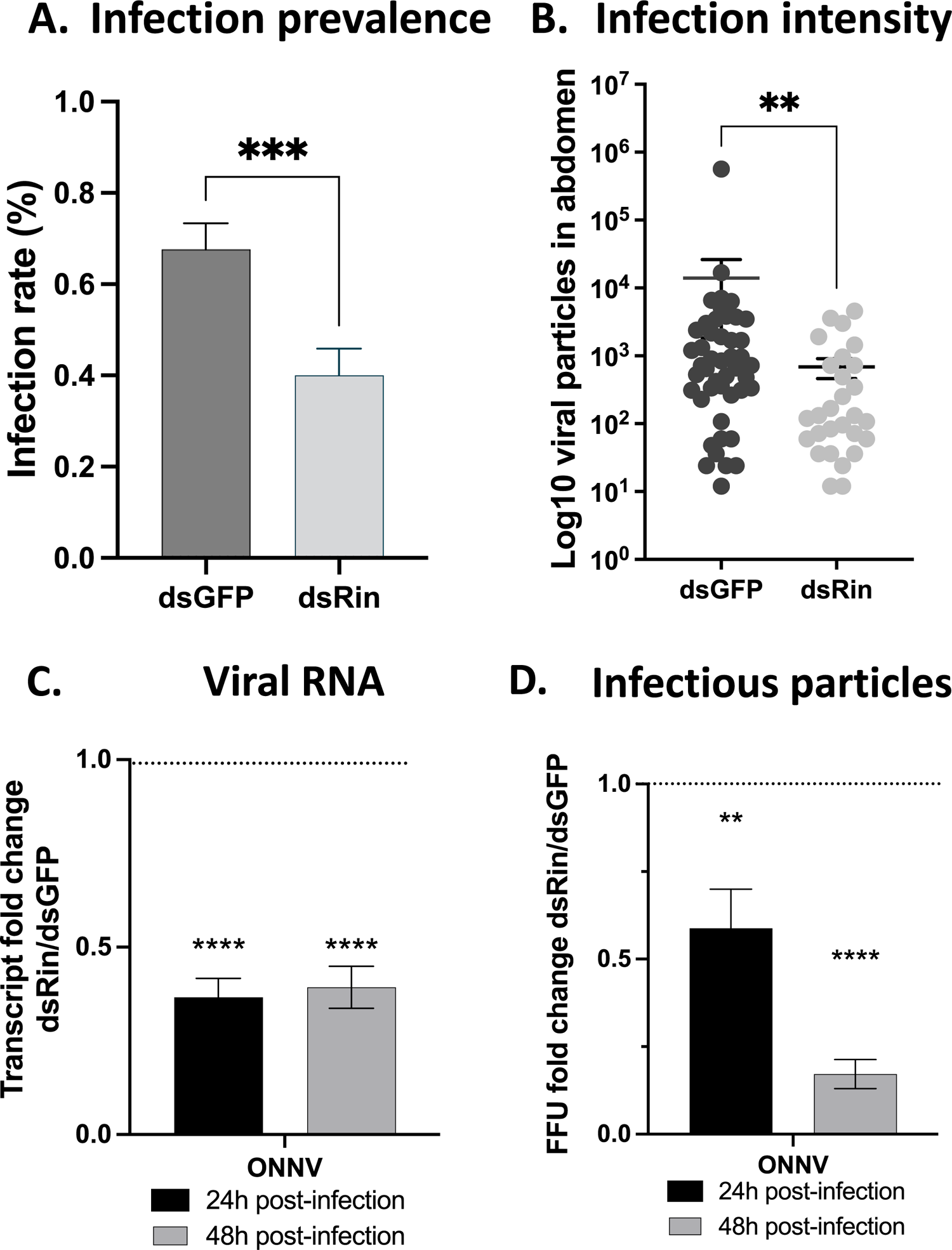
Rin is a proviral factor for ONNV infection in Anopheles. **A, B**. An. coluzzii mosquitoes after bloodmeal infection. **C, D**. An. coluzzii 4a3A hemocyte-like cells. **A, B**. Female mosquitoes were treated with dsRNA two days prior to an infectious bloodmeal. On the day of infection, the unfed females were removed and used to test Rin silencing efficiency, then on 6 d post-bloodmeal ONNV infection in abdomens was measured by viral titration. **A**. The effect of Rin depletion on infection prevalence among bloodfed mosquitoes as a proportion of the total mosquitoes analyzed. y-axis, percentage of abdomens positive for ONNV in dsRin or dsGFP-treated mosquitoes. **B**. The effect of Rin depletion on infection intensity in the positive mosquito samples. y-axis, load of viral particles, each point represents the viral titer in one abdomen. Error bars indicate the SEM. Results from 3 independent experiments with 72 total mosquitoes analyzed per condition. ** P < 0.01, *** P < 0.005, ns non-significant. (C-D) Rin is a proviral factor for ONNV infection in 4a3A An. gambiae cells. Rin proviral actions on viral RNA quantity (C) and infectious particle production (D) are showed. (C-D) Each bar represents the level of transcript abundance of ONNV RNA (C) or focis forming unit (D) relative to dsGFP-treated control (defined as 1.0), and error bars indicate the SEM. Histogram showed are the results of three independent experiments. *** P < 0.005, **** P < 0.0001.

**Figure S2.**
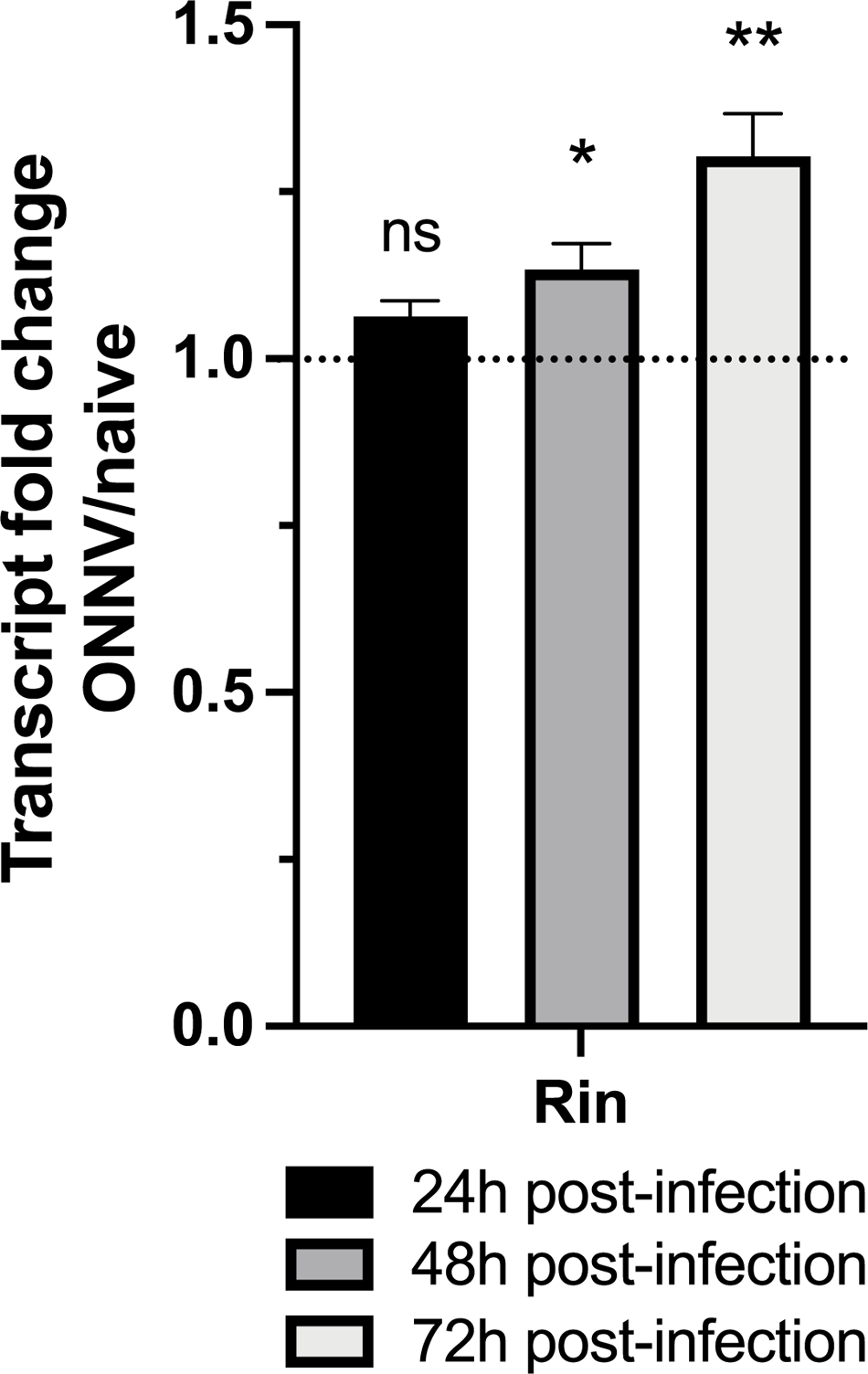
Rin transcriptional expression is induced by ONNV in 4a3A An. gambiae cells. (A-B) Rin transcriptional expression (A) during ONNV infection is showed. (A) Each bar represents the level of transcript abundance of Rin in ONNNV-infected cells relative to naïve cells (defined as 1.0), and error bars indicate the SEM. Histogram showed are the results of three independent experiments. * P < 0.05, ** P < 0.01, ns non-significant.

**Figure S3.**
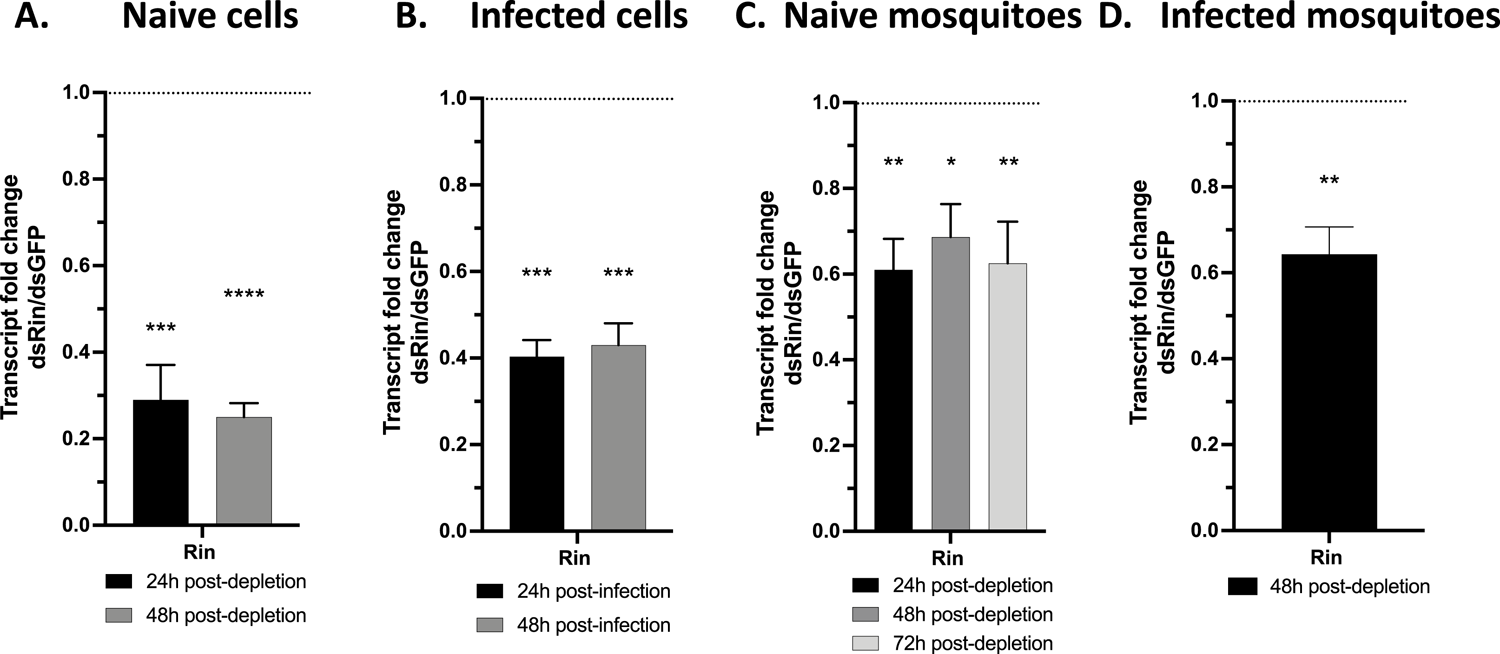
Rin transcript is efficiently depleted by the dsRNA treatment in both 4a3A An. coluzzii in vitro model and in An. coluzzii mosquito in vivo model. (A-B) Rin transcript after dsRin treatment in naïve cells (A) or infected cells (B) are showed. (C-D) Rin transcript after dsRin treatment in naïve mosquitoes (C) or the day of an infectious blood feeding (D) are showed. (A-D) Each bar represents the level of transcript abundance of Rin relative to dsGFP-treated control (defined as 1.0), and error bars indicate the SEM. Histogram showed are the results of at least three independent experiments. Results on mosquitoes are analyzed from a pull of at least 6 mosquitoes. *P < 0.05, ** P < 0.01, *** P < 0.005, **** P < 0.0001, ns non-significant.

**Figure S4.**
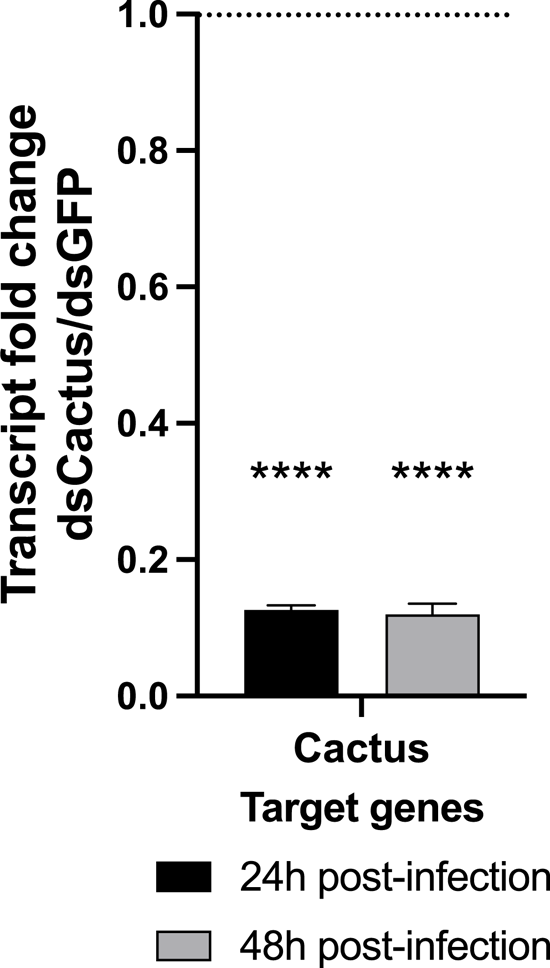
Rin, Rel1 and Cactus transcripts are efficiently depleted by the dsRNA treatment in 4a3A An. gambiae in vitro model. (A-B) Rin and Rel1 transcript after dsRNA treatment in infected cells at 24h (A) and 48h (B) post-infection are showed. (C) Cactus transcript after dsCactus treatment in infected cells is showed. (A-C) Each bar represents the level of transcript abundance of Rin, Rel1 or Cactus relative to dsGFP-treated control (defined as 1.0), and error bars indicate the SEM. Histogram showed are the results of at least three independent experiments. *P < 0.05, ** P < 0.01, *** P < 0.005, **** P < 0.0001.

**Figure S5.**
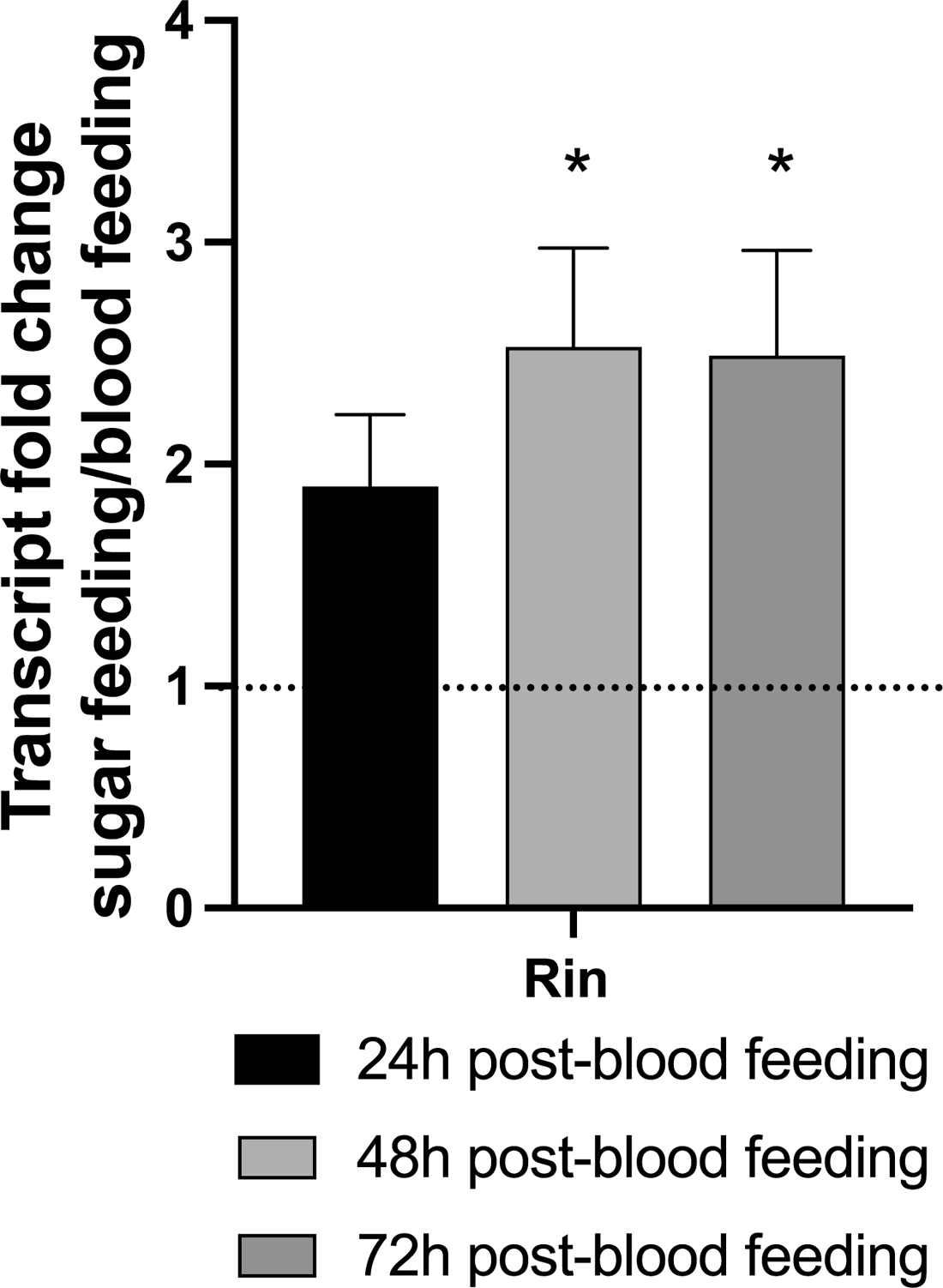
Rin transcript is enhanced after the blood-feeding of An. coluzzii mosquitoes. Rin transcript level in blood-fed female mosquitoes is showed. Each bar represents the level of transcript abundance of Rin in blood-fed mosquitoes relative to sugar-fed mosquitoes (defined as 1.0), and error bars indicate the SEM. Histogram showed are the results of at least three independent experiments. A pool of at least 6 mosquitoes has been analyzed by biological replicates. *P < 0.05.

